# The possible role of lipid bilayer properties in the evolutionary disappearance of betaine lipids in seed plants

**DOI:** 10.1101/2023.01.24.525350

**Authors:** Stéphanie Bolik, Alexander Schlaich, Tetiana Mukhina, Alberto Amato, Olivier Bastien, Emanuel Schneck, Bruno Demé, Juliette Jouhet

## Abstract

Phosphate is vital for plant and algae growth, yield, and survival, but in most environments, it is poorly available. To cope with phosphate starvation, photosynthetic organisms used their phospholipids as a phosphate reserve. In microalgae, betaine lipids replace phospholipids whereas, in higher plants, betaine lipid synthesis is lost, driving plants to other strategies. The aim of this work was to evaluate to what extent betaine lipids and PC lipids share physicochemical properties and could thus substitute each other. Using neutron diffraction and molecular dynamics simulations of two synthetic lipids, dipalmitoylphosphatidylcholine (DPPC) and dipalmitoyl-diacylglyceryl-N,N,N-trimethylhomoserine (DP-DGTS), we show that DP-DGTS bilayers are thicker, more rigid, and mutually more repulsive than DPPC bilayers. The different properties and hydration response of PC and DGTS provide an explanation for the diversity of betaine lipids observed in marine organisms and for their disappearance in seed plants.

## INTRODUCTION

Many organisms, such as bacteria, plants, and fungi, rely on mineral nutrients taken up directly from the environment, and therefore specialize their metabolism to cope with their limited availability. For example, most organisms have well-defined responses to phosphate limitation, including the replacement of cellular membrane phospholipids with non-phosphorous lipids. This phenomenon has been well documented in plants [1,2], where glycolipids replace phospholipids; whereas in bacteria, fungi or microalgae, membrane phospholipids could also be replaced by another kind of non-phosphorous lipids called betaine lipids [3–6]. Presently, three basic types of betaine lipids are known in photosynthetic organisms; DGTS (diacylglyceryl-N,N,N-trimethylhomoserine), DGTA (diacylglycerohydroxymethyl-N,N,N-trimethyl-b-alanine), and DGCC (diacylglycerylcarboxy-N-hydroxymethylcholine) [7–9]. They are diversely represented in organisms, DGTS being the most frequent whereas DGTA and DGCC have been solely observed in marine organisms[10]. In algae, under phosphate starvation, a situation commonly met in the environment, betaine lipids replace phospholipids in extraplastidic membranes. Because betaine lipids are localized in these membranes [11,12] and share a common structural fragment with the main extraplastidic phospholipid phosphatidylcholine (PC) (Figure 1A and B), it can be speculated that these two lipid classes are interchangeable, but this was never demonstrated. Betaine lipid synthesis is located in the ER [13,14] and betaine lipids are expected to be absent in photosynthetic membranes [12]. Therefore, this PC-betaine lipid replacement is not expected to occur in photosynthetic membranes. However, it might occur at the surface of the chloroplast envelope where PC might be present [15–17]. Nothing is known about the composition of mitochondrial membranes in algae but because PC is a major lipid component in plant and fungal mitochondria, this replacement might also occur in mitochondria.

**Figure 1.**
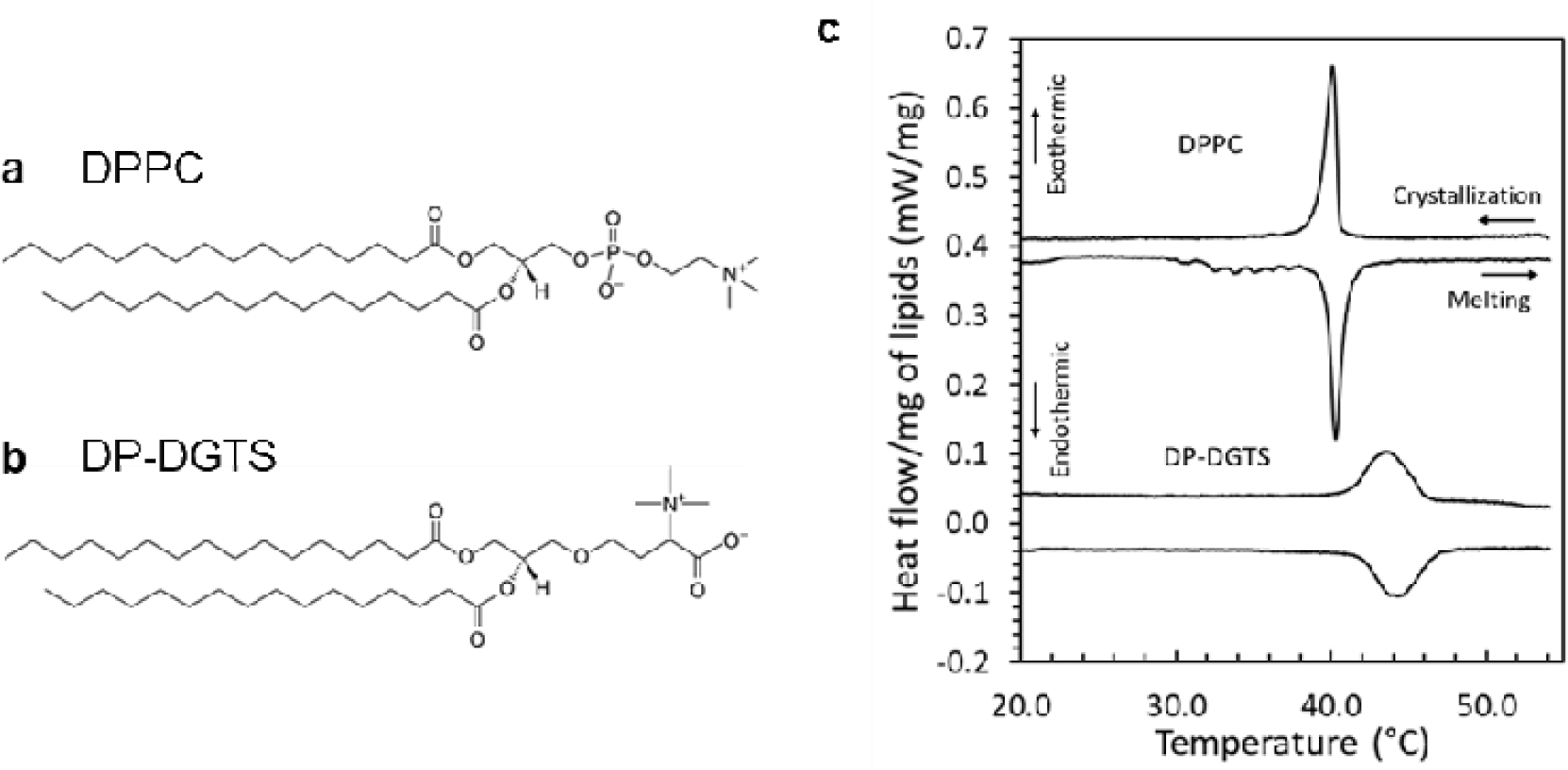
Study of the phase transition of DPPC and DP-DGTS. **a and b**. Lipid chemical structures of (a) Dipalmitoylphosphatidylcholine (DPPC) and **(b)** Dipalmitoylglyceryl-N,N,N-trimethyl-homoserine (DP-DGTS). **c.** Differential Scanning Calorimetry (DSC) thermograms of DPPC and DP-DGTS unilamellar vesicles. DPPC curves were vertically offset by 0.4 mW/mg for clarity.

The identification and description of betaine lipids in cells are so far only based on biochemical analyses and therefore poorly cover the wide range of organisms [7,10,18–20]. In a simplified view, betaine lipids are present in non-vascular plants, algae and some fungi, but even these groups show several exceptions, and they are absent from seed plants, i.e. gymnosperms and angiosperms [21]. The presence of betaine lipids is not linked to the synthesis of betaine, a soluble compound present in almost every organisms including most animals, plants, and microorganisms, acting as protectant against osmotic stress [22]. All these studies were accomplished before the amendment of the classification and the use of genetic tools to improve the phylogeny. Because the biosynthetic pathway to DGTS is now known, new phylogenetic inference on the appearance and disappearance of DGTS biosynthesis enzymes could be achieved.

Biosynthesis of DGTS was first discovered in bacteria and is realized in two steps by the two enzymes BtaA and BtaB whereas in *Chlamydomonas* and in fungi it is catalyzed by one bifunctional enzyme Bta1 containing the BtaA and BtaB domains. BtaA uses diacylglycerol and S-adenosylmethionine (SAM) as substrates to produce diacylglycerol-O-homoserine that will then be used by BtaB to produce DGTS by adding successively three methyl groups on the nitrogen from the SAM donor [5,14]. DGTA is assumed to be synthesized by decarboxylation/recarboxylation of DGTS [23] but the enzyme and the mechanisms are still unknown and nothing is known so far about DGCC synthesis. The reconstruction of phylogenetic relationships could help to decipher the evolution of the lipid metabolism from algae to higher plants, the adaptation of plant and algae strategy to nutrient starvation and the disappearance of betaine lipid synthesis pathways through evolution.

To unravel the functions of betaine lipids as well as their ability to act as phosphorous-free PC lipid substitutes, we investigated their physicochemical properties. The only physicochemical study was carried out by Sato and Murata in 1991 [24]. They showed that the gel-to-fluid phase transition temperature of DGTS bilayers is only slightly higher than that of PC lipids, suggesting similarities in the headgroups’ mutual interactions. To which extent betaine lipids are suited to replace PC lipids also depends on other factors such as structural aspects of lipid packing and their influence on membrane fluidity, bending rigidity, and on membrane-membrane interactions. Here, we have investigated these properties by neutron diffraction and computer simulations.

## RESULTS

### Phase transition for DP-DGTS bilayers is broader than for DPPC bilayers

Lipids with the same diacylglycerol backbone were chosen to study molecules that differ only by their polar head. Because DPPC has been widely studied and DP-DGTS is commercially available, our work was achieved with these two molecules with saturated C16 tails (Figure 1 A and B). To determine the chain melting temperature for DPPC and DP-DGTS, differential scanning calorimetry (DSC) thermograms were collected from monodisperse unilamellar vesicles obtained by extrusion (Figure 1C).

The DSC data show a sharp phase transition at 40.2 ± 0.1°C for DPPC corresponding to the transition between the ripple phase and the fluid phase, which is consistent with earlier reports on DPPC large unilamellar vesicles [25]. For DP-DGTS, the phase transition is much broader and occurs within a range of 4°C around 43.8°C, which confirms previous report [24]. With this in mind and to favor the fluid phase state, all following experiments were carried out at 50°C, at least 5°C above the phase transition in excess water.

We then reconstituted multilayer stacks of membranes from pure DPPC or DP-DGTS at the surface of silicon wafers and collected neutron diffraction patterns (Figure S1) that contained three or four diffraction orders, indicating that the lattice disorder of the sample is small [26]. In the studied range of relative humidities (i.e., covering high to low osmotic pressures), the coexistence of gel and fluid phase was observed with two sets of Bragg peaks (Figure S1E and E’, Figure S2), each set corresponding to a lamellar phase, the gel phase to the peaks at wider angles and the fluid phase to the peaks at smaller angles.

**Fig. S1.**
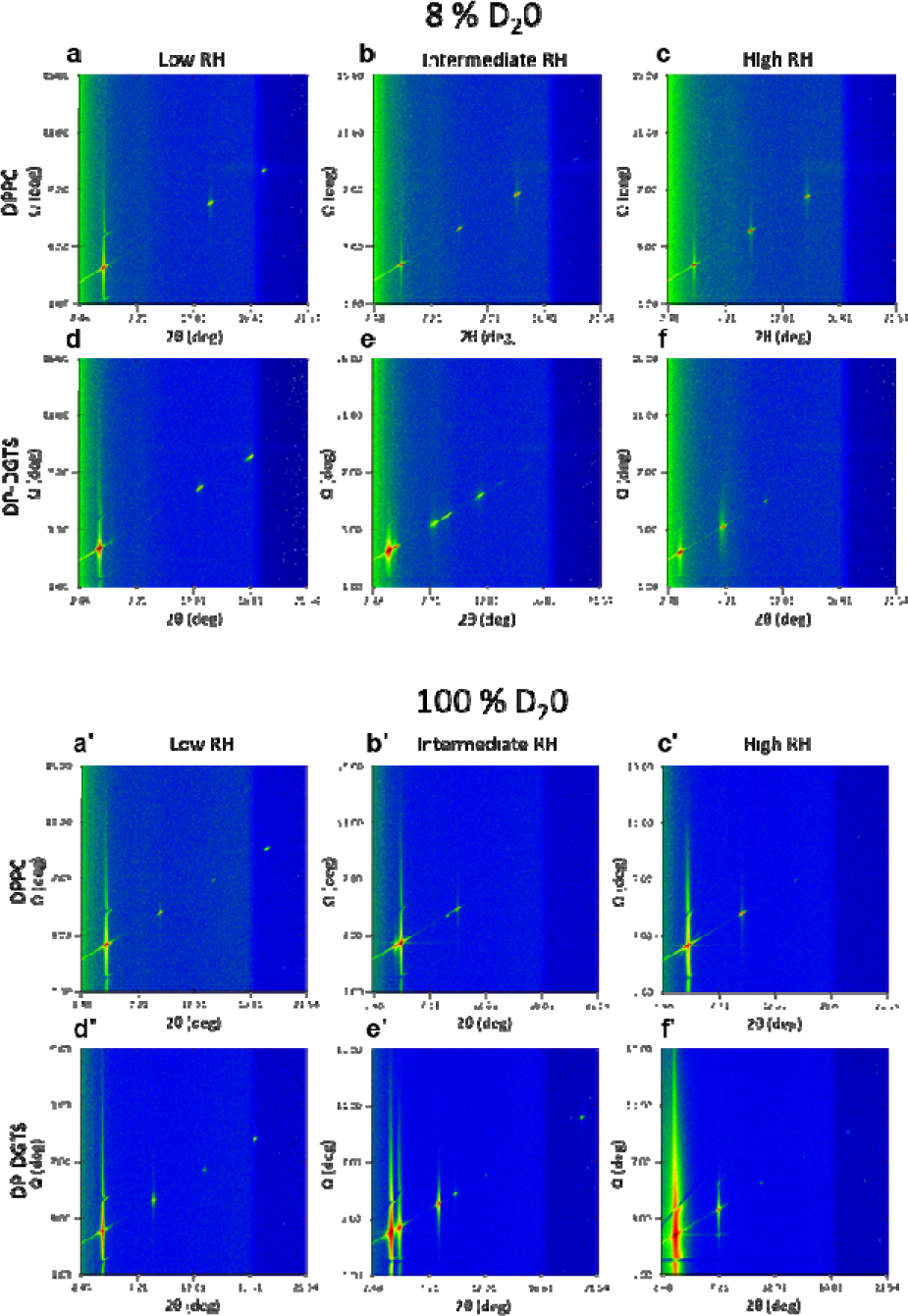
Diffraction scan I(2*θ*, Ω) showing the Bragg peak positions of DPPC (**a-c** and’ **a**’-**c**’) and DP-DGTS (**d-f** and **d**’-**f**’) at three different humidities: low (**a, a**’ and **d, d**’) intermediate (**b, b**’ and **e, e**’) and high (**c, c**’ and **f**, f’). The measurements were done at 50°C. a to f: Solvent composition 8% D_2_O. Relative humidity values: DPPC 30.3, 85.0, 95.8%. DP-DGTS 32.2, 87.2, 97.9%. **a**’ **to f**’: Solvent composition 100 % D_2_O. Relative humidity values: DPPC 30.3, 85.0, 95.8%. DP-DGTS 32.2, 87.2, 97.9%.

**Fig. S2.**
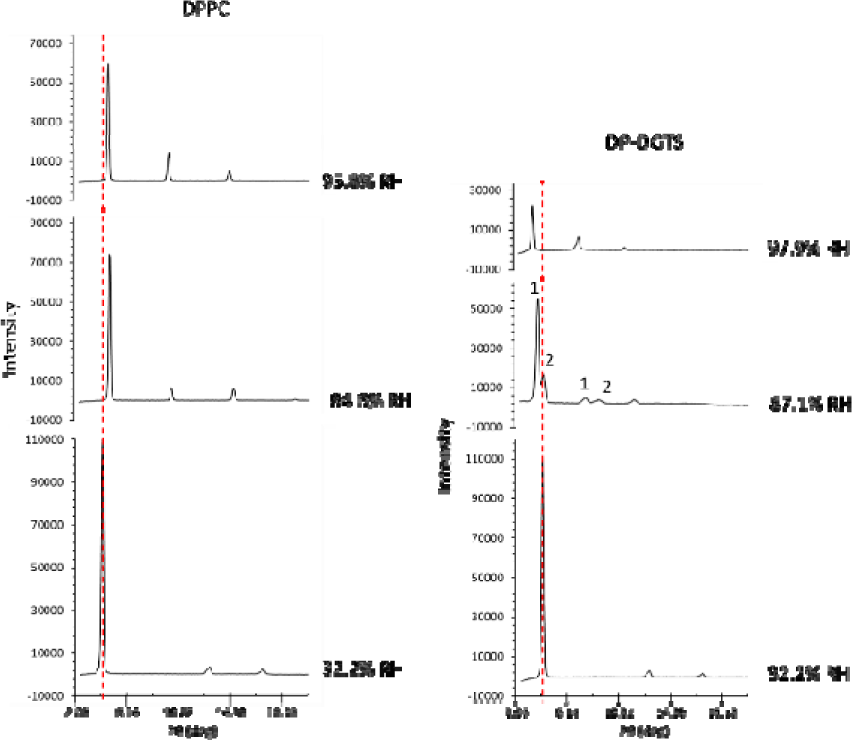
Intensity of the Bragg peaks at 3 humidities. The gel and fluid phases in the DP-DGTS sample at RH=87.1% are noted 1 and 2. Phase 1 is probably in the gel state and phase 2 the fluid one. The temperature of the sample was 50°C.

As described in previous studies [27–29], we extracted from these data the pressure-distance curves, i.e., the dehydrating osmotic pressure Π as a function of the repeat distance d. As seen in Figure 2A, by increasing the humidity, DPPC molecules transit from the gel to the fluid phase via a ripple phase through a narrow window of osmotic pressures as previously reported [30,31]. In contrast, DP-DGTS bilayers show a phase coexistence that can be observed over a wide Π-range and without the appareance of a third phase that could be attributed to a distinct ripple phase (Figure 2B) before forming a single fluid phase at high humidity (i.e., at low Π). Based on DSC and neutron diffraction as two independent techniques, we can safely conclude that the phase transition for DP-DGTS is broad. This observation indicates that the free energy difference between the two phases is very small over a wide osmotic pressure range and may be connected to the shapes of the pressure-distance relations in the two phases, which are discussed further below.

**Figure 2.**
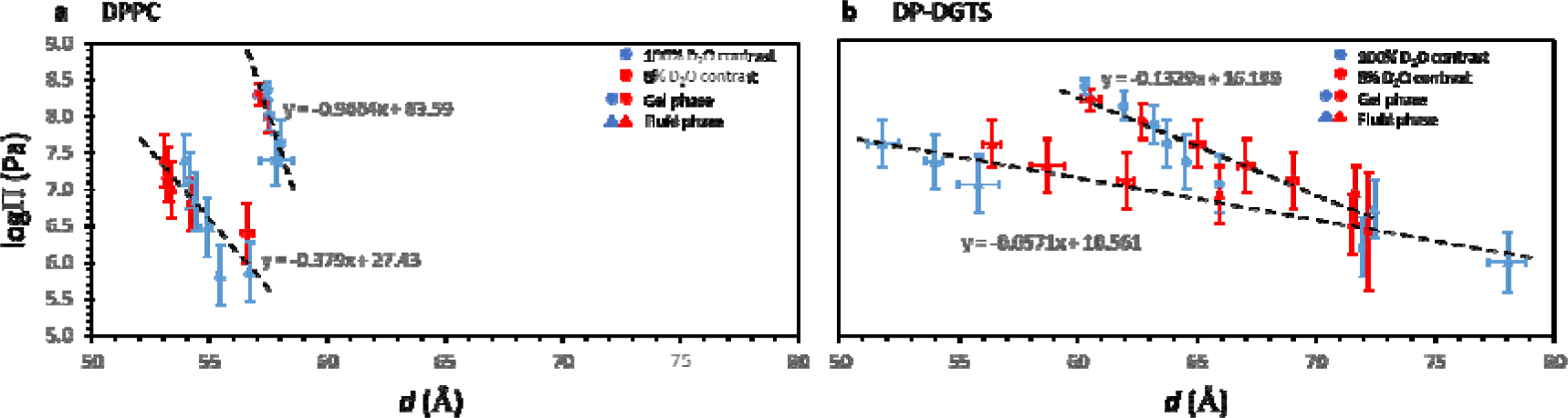
Lamellar period d of DPPC (**a**) and DP-DGTS (**b**) solid supported multilayers as a function of the applied osmotic pressure Π, measured at 50°C. The period was determined by neutron diffraction at 100% D_2_O contrast (blue), and at 8% D_2_O contrast (red). The coexistence between gel and fluid phase is shown with the circles and the triangles respectively.

### Hydration repulsion is stronger between DP-DGTS bilayer than between DPPC bilayers

Hydration forces are repulsive and of short range [32]; they are important for diverse processes such as preventing membrane fusion and adhesion [33] or protein adsorption [34]. To evaluate hydration forces, using membrane diffraction data at 8 % D_2_O contrast as described in material and method and Figure S3, we extracted the water layer thickness d_w_ for each osmotic pressure (Figure 3). The exponential fit to the data, in Figure 3A and B, yields the decay length λ_w_, representing the short-range repulsion between the bilayers, which is dominated by hydration forces [35].

**Fig S3.**
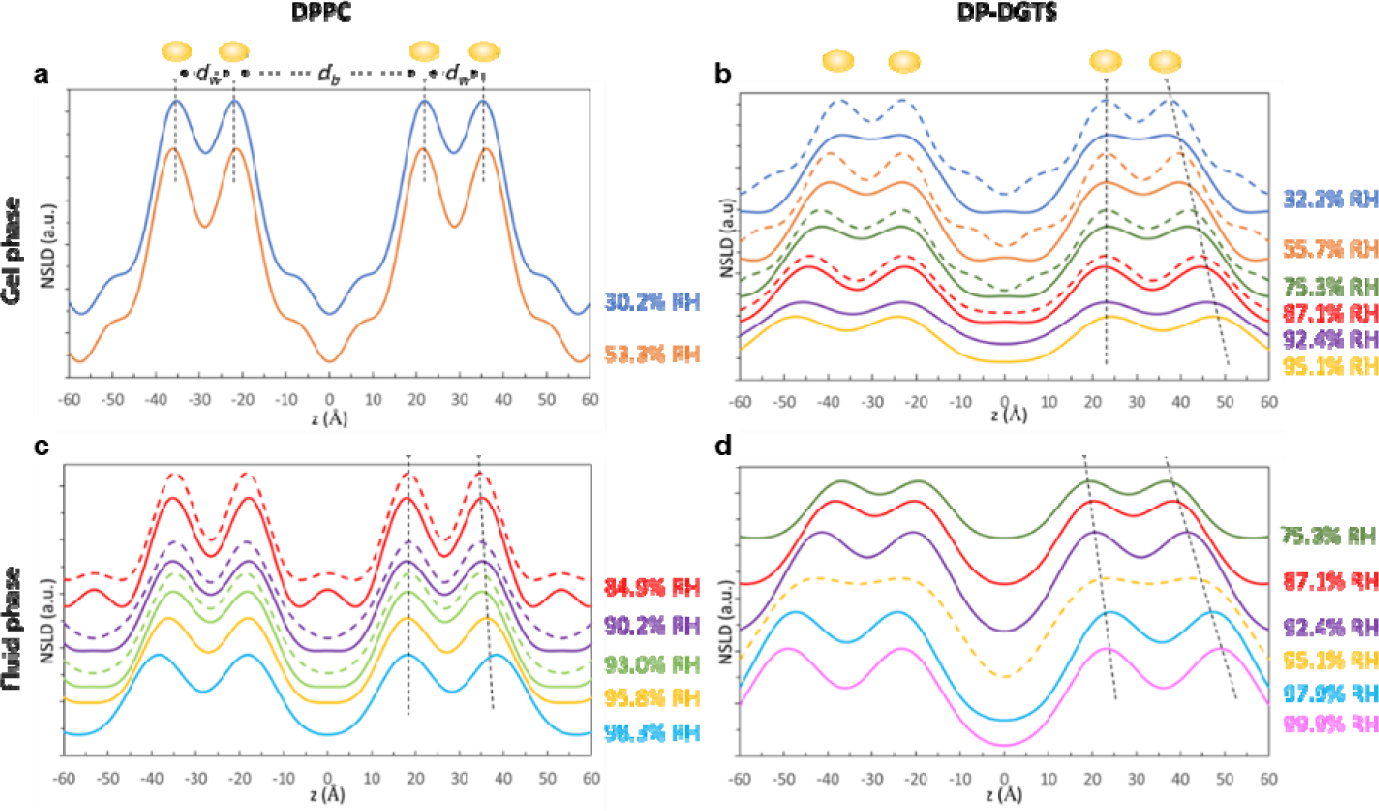
Neutron Scattering Length Density (NSLD) of DPPC (**a** and **c**) and DP-DGTS (**b** and **d**) measured at 50°C as described in [27]. The gel and fluid phase are separated in two graphs for the two lipids, **a** and **b** for the gel phase, **c** and **d** for the fluid phase. The bilayer thickness d_b_ corresponds to the center to center distance between headgroups as described in [29] and the water layer thickness d_w_ corresponds to d-d_b_ with d being the lamellar repeat distance or d-spacing and d_b_ the bilayer thickness. These are represented in panel **a.** The solid lines are the NSLD profile calculated using the minimum of three Bragg peaks and the dash lines are calculated using 4 or 5 peaks when visible at the highest humidities for DP-DGTS. The dotted black lines show the shift upon hydration of the maxima in the SLD profile, showing the increase of the water layer thickness.

**Figure 3:**
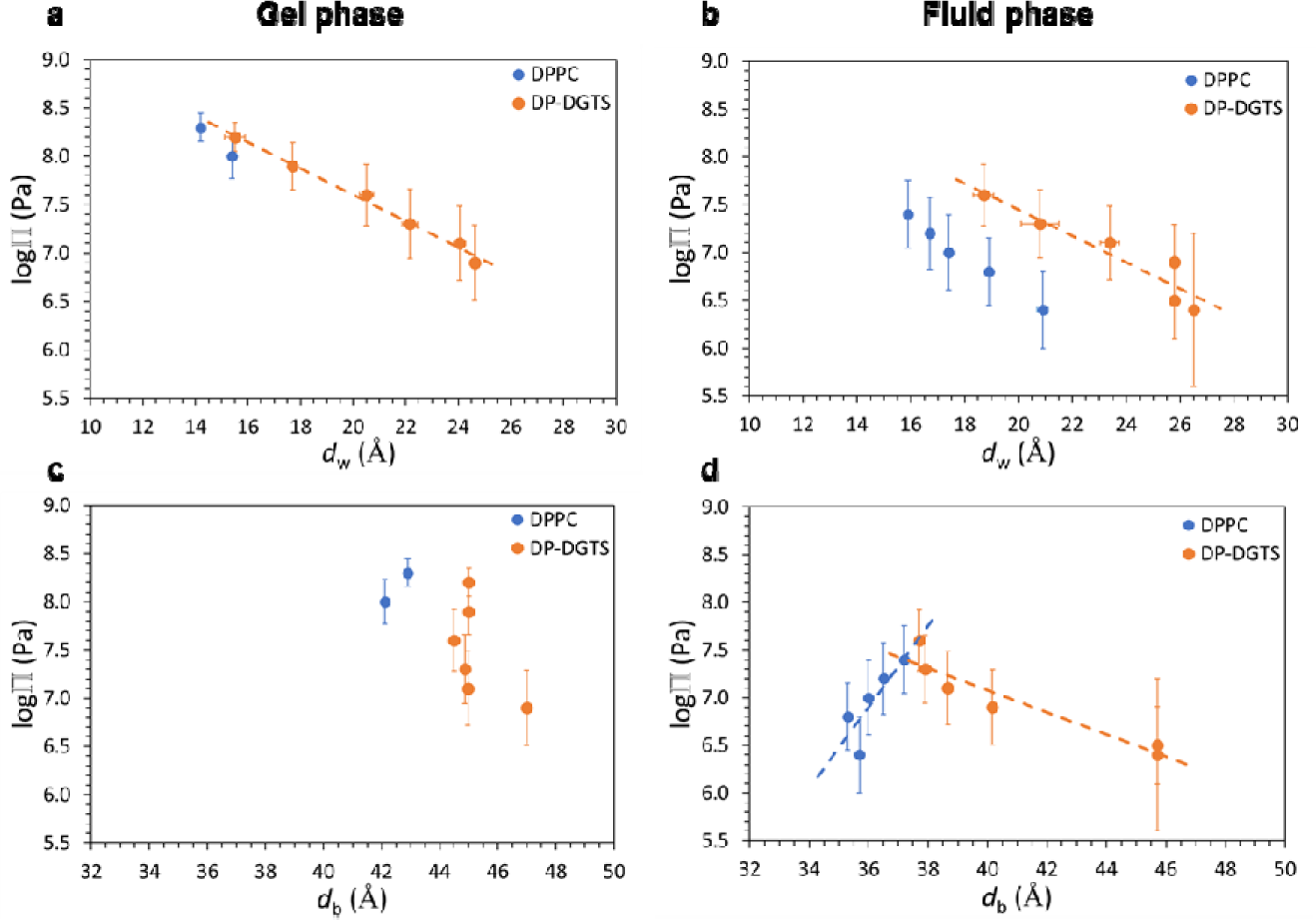
**(a and b)** Pressure-distance (d_w_) and (**c and d**) pressure-bilayer thickness (d_b_) curves of DPPC (blue) and DP-DGTS (orange), measured at 50°C. The gel phase is shown on graphs A and C, and the fluid phase on graphs B and D.

For the gel phase of DPPC, because we only have two data points, we could not evaluate the decay length. In the fluid phase, we found for DPPC λ_w_^(fluid)^ = 5.2 Å, in acceptable agreement with the 4 Å reported by Kowalik et al.[35]. For DP-DGTS, because the gel and fluid phases coexist in a wide range of pressures, we could calculate the decay length of both. We obtain λ_w_^(gel)^ = 6.9 Å and λ_w_^(fluid)^ = 7.7 Å. λ_w_^(gel)^ is about 3 times higher for DP-DGTS bilayers than what is reported in the literature for DPPC (λ_w_^(gel)^ = 2.1 Å [35] whereas λ_w_^(fluid)^ is only 1.4 times higher for DP-DGTS bilayers than for DPPC. Moreover, there is no significant difference in the decay length between gel and fluid phases in DP-DGTS bilayers, which is in contrast to DPPC. These differences suggest that the hydration repulsion between the surfaces of these two bilayers are significantly different.

Short-range interactions between adjacent bilayers in a stack could be represented by the inter-membrane compression modulus B [29]. As explained in the materials and methods section and supported by Figure S4, we could extract B=29 MPa for DPPC at RH=99%. This is consistent with an earlier report of B=46 MPa measured at 60°C and 95 % RH [36], because B decreases systematically with increasing hydration due to the associated decay of the membrane interactions[37]. For DP-DGTS, B=1.5 MPa, a much smaller value than the one for DPPC. The difference can be attributed to the water layer, which is thicker for DP-DGTS at this humidity (Figure 3B). Therefore, the compression modulus of DP-DGTS is lower than that of DPPC, reflecting weaker interactions between DP-DGTS bilayers than between DPPC bilayers.

**Fig. S4:**
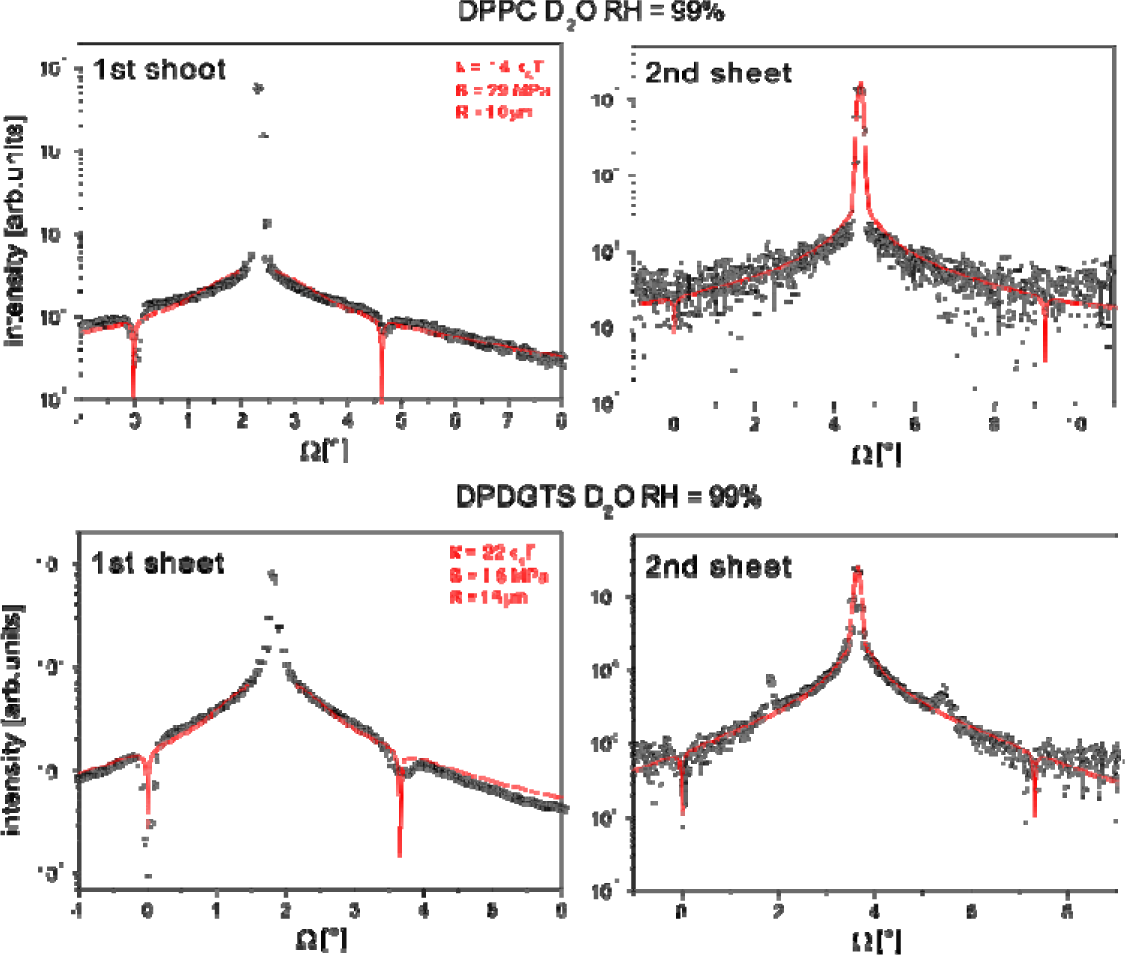
2q-integrated Bragg sheet intensities of DPPC membranes (top) and DP-DGTS membranes (bottom) at RH=99 % of the first (left) and second (right) Bragg sheets as a function of Ω, featuring the respective central specular maxima symmetrically flanked by the slowly decaying diffuse scattering intensity. The latter is locally decorated with minima at conditions of high absorption (Ω ≈ 0 and Ω ≈ 2q) and peaks arising from multiple scattering effects [38,39]. The solid lines (in red) superimposed to the experimental data points represent simulated Bragg sheet intensities corresponding to the best-matching parameters in the continuum-mechanical model simultaneously describing the first and second Bragg sheets as explained in the material and method section. Absorption close to Ω ≈ 0 and Ω ≈ 2q was modeled as described in [38]. The best-matching model parameters for both systems are summarized with the empirical cut-off parameter termed R and the mechanical parameters k and B derived thereof according to Eqs. 6 and 7 as described in the material and method section [39].

To evaluate the long-range interaction between lipid bilayers, we determine the lamellar period of the bilayer at full hydration, by studying multilamellar vesicles (MLV) of DPPC and DP-DGTS in excess water by Small Angle Neutron Scattering (SANS) (Figure 4). For DPPC, the periodic lamellar structure indicates a repeat distance of 65.8 Å in the gel phase at T=20°C and 67.8 Å in the fluid phase at T=50°C, confirming the literature values [40]. However, for DP-DGTS a Bragg peak could hardly be identified, indicating an only very weak correlation between membranes. A fit to the weak intensity modulations yields the repeat distance of 190 Å in the gel phase at 20°C and 210 Å in the fluid phase at 50°C.

**Figure 4:**
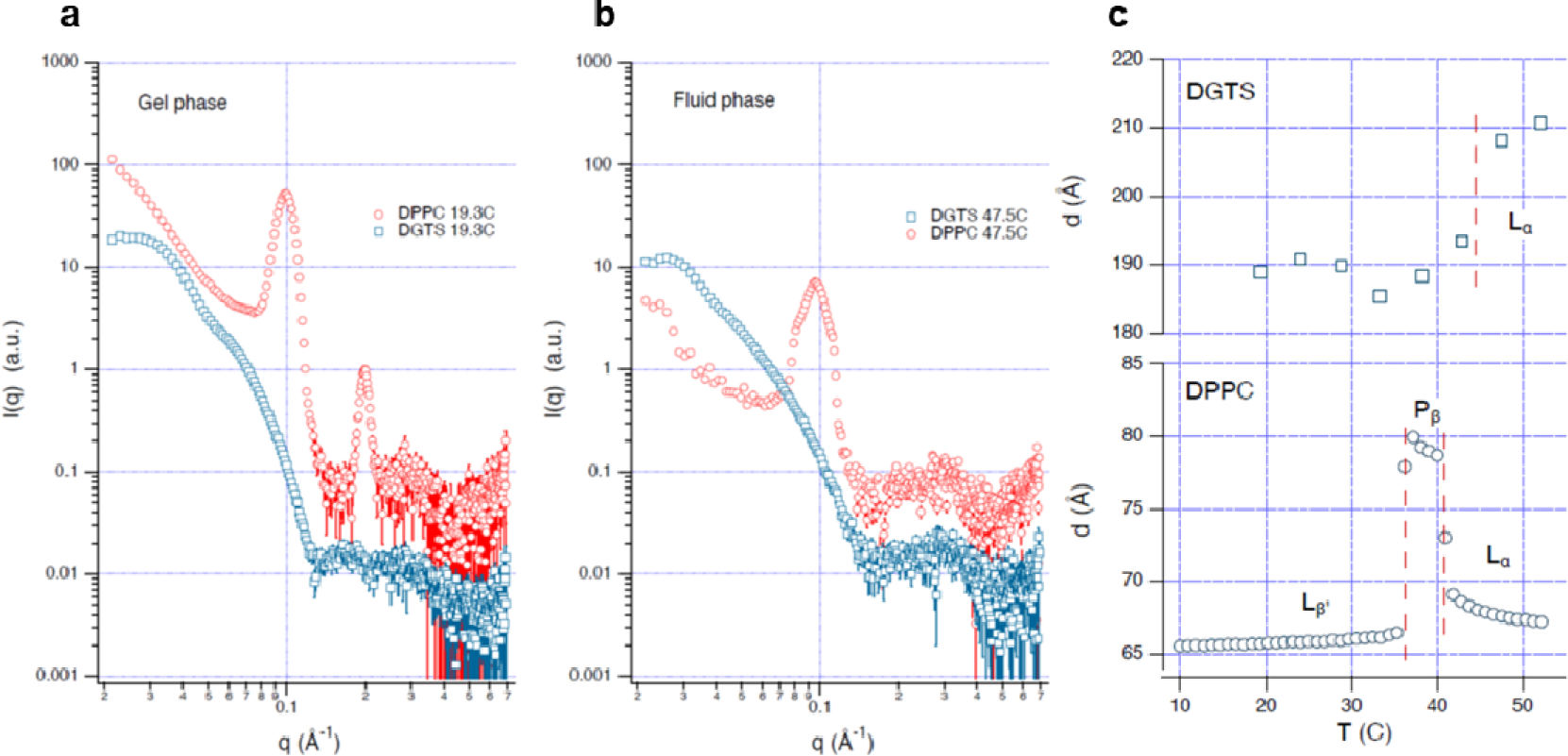
**a and b**: SANS curves of multilamellar vesicles in excess water made of DP-DGTS (blue symbols, scaled by 10) and DPPC lipids (red symbols) in gel phase **a** (T = 19.3°C) or in fluid phase **b** (T= 47.5°C). **c**: Lamellar period of DP-DGTS and DPPC as a function of temperature. For DPPC the two phase transition temperatures are directly deduced from the sharp period changes and represented by the vertical dashed lines. For DP-DGTS, where no sharp change is observed the dashed line corresponds to the gel-fluid transition temperature (43.8°C) obtained by DSC (Figure 1). No ripple phase P*_β_* was detected for DP-DGTS bilayers.

The behavior of DP-DGTS indicates the presence of a long-range repulsive contribution to the interaction between DP-DGTS bilayers that is not present for DPPC bilayers. On top of the hydration repulsion, the repulsion may be strengthened due to the presence of charges. Namely, one cannot exclude that a small fraction of lipids deviates from the zwitterionic state, resulting in a tiny effective surface charge, which might be more pronounced for DP-DGTS because the pKa values of phosphates and carboxyl groups are not identical. Altogether, these data indicate that the interaction between DP-DGTS bilayers is more repulsive than that between DPPC bilayers, both at short and long-range.

**DP-DGTS bilayers are thicker and more rigid than DPPC bilayers.**

Because the bilayer thickness affects membrane protein insertion as well as membrane mechanical properties, we also analyzed the bilayer thickness d_b_ as a function of Π (Figure 3C and D). The d_b_ values obtained for DPPC are 42.5 ± 0.6 Å and 36.1 ± 0.7 Å respectively for the gel and the fluid phase, with a slight decrease in d_b_ when Π increased. These values are slightly smaller than those described in the literature (44.2 Å for the gel phase [29] and 37.2 Å for the fluid phase [41], consistent with our bilayer thickness definition, measured between the middle of the two opposite polar heads and therefore not comprising the complete polar heads as the references cited above. Our SLD analysis from Figure S3 and Figure 3C and D reproduces the known dehydration-thickening of fluid PC lipid multi-bilayers. Indeed, the slight increase of the bilayer thickness upon osmotic dehydration is expected because the hydration favors disordering of the lipid components of the bilayer as previously reported for DOPC [42]. This phenomenon explains the transition between the gel and the fluid phase by simply decreasing Π at constant temperature. Altogether, by comparison with literature, the results obtained on DPPC bilayers validate the methodology employed here.

By following the same experimental procedure as for DPPC bilayers, we observed for DP-DGTS a decrease of bilayer thickness between the gel and fluid phase, as in DPPC bilayers and as expected upon fatty acid chain melting. However, whatever the phase, the thickness of DP-DGTS bilayers (45.2 ± 0.9 Å in gel phase and 40.9 ± 3.8 Å in fluid phase) is significantly higher than that of DPPC. Furthermore, the DP-DGTS bilayer thickness in fluid phase seems to respond differently to hydration. In fact, whereas DPPC bilayers follow a dehydration-thickening trend, DP-DGTS bilayers seem to follow a dehydration-thinning trend.

The membrane bending modulus κ is a measure of the energy required to bend a membrane from planarity to a defined curvature. This value, as the compression modulus B mentioned earlier, is accessible by Bragg sheet analysis (Figure S4 and materials and methods). We obtained κ ≈ 14 ± 2 k_B_T and κ ≈ 22 ± 2 k_B_T for DPPC and DP-DGTS, respectively. The value found for DPPC is in agreement with the literature (18.3 ± 3.1 k_B_T or 15.0 ± 1.6 k_B_T) [43]. The higher bending modulus for DP-DGTS can be explained with the higher bilayer thickness. In fact, according to classical beam theory the bending rigidity of a homogeneous planar object scales with the third power of the thickness. Application of this relation to the bilayers while simplifying them as homogeneous objects yields the

estimate 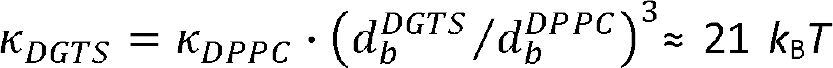, which is in good agreement with our data on bilayer thicknesses.

### Molecular dynamics simulations confirm stronger hydration repulsion of DGTS and reveal differences in the water distribution

To further investigate the differences observed between DPPC and DP-DGTS bilayers, we employed atomistic molecular dynamics (MD) simulations. The Thermodynamic Extrapolation method [44,45] allows us to perform simulations at prescribed water chemical potential and thus to obtain the interaction pressures between membranes with high precision (Figure S5A and B). Snapshots of our simulations in fluid and gel states are presented in Figure S6.

**Fig S5:**
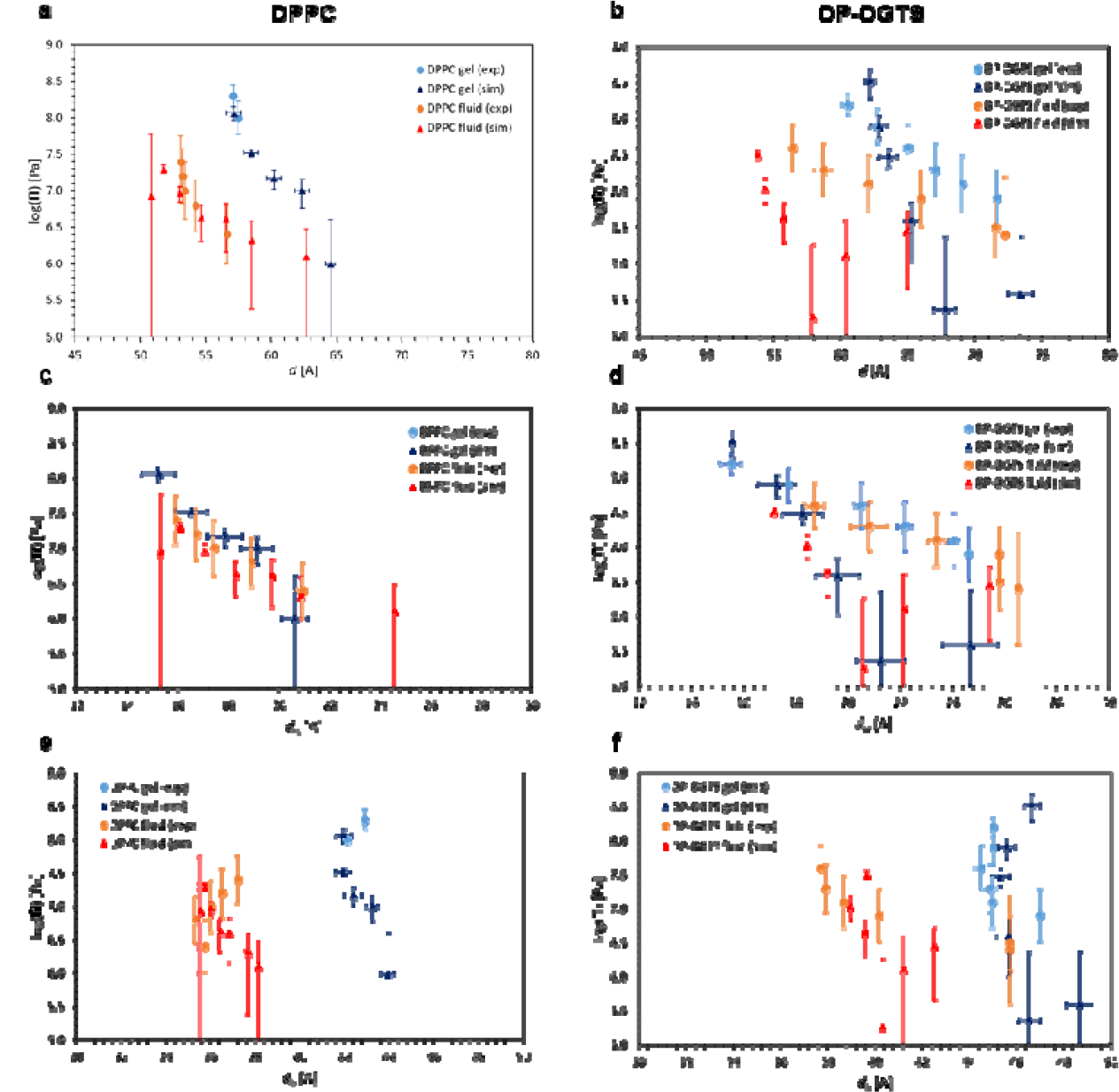
**a and b**. Simulated lamellar period of DPPC (**a**) and DP-DGTS (**b**) bilayer stacks as a function of the humidity, measured at 50°C (circle) or computed (triangle). **c to f**. Comparison of the bilayer thickness (**c and d**) and of the water layer thickness (**e and f**) as a function of the hydration pressure obtained by simulation (triangles) or by experiment (circles) in DPPC bilayers (**c and e**) and in DP-DGTS bilayers (**d and f**).

**Fig S6:**
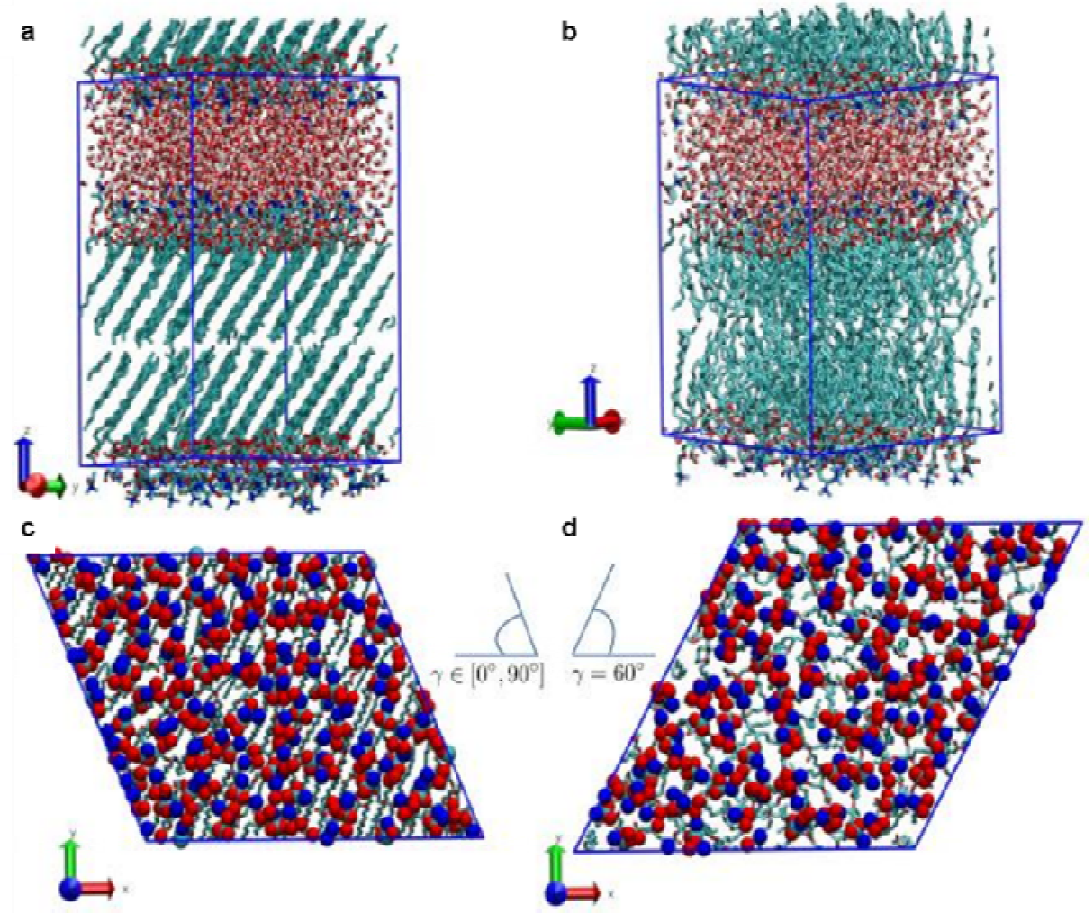
**a,b**. Snapshots of the simulations in the L_β_ (gel) and L_α_ (fluid) phases, respectively. Blue lines show the periodic simulation box. **c.** Top-view of a simulation in the gel phase where the head-group atoms are highlighted using spheres. The tilt angle is free to adjust according to the lipid lattice structure with a typical value of. **d.** For the simulations in the fluid phase the tilt angle is fixed at. Note that due to the minimum image convention in the periodic boundary conditions employed, i.e. the box angle in **C** is allowed to flip its direction.

We extracted from the simulations the Π dependence of the bilayer thickness (Figure S5C and D) and of the water layer thickness (Figure S5E and F) and compared the experimental data (circles) to the simulations (triangles). While the simulations largely reproduce the experimental data for DPPC, indicating that the simulation model represents well the hydration forces between DPPC bilayers, the hydration dependence of the bilayer thickness of DP-DGTS is not so well reproduced. However, the simulations clearly confirm that DP-DGTS bilayers are thicker and interact more repulsively than DPPC bilayers, although the difference obtained by the simulations is not quite as pronounced as observed in the experiments. This discrepancy may be attributed to the fact that any deviation from charge-neutrality, which may occur in the experimental system due to lipid (de-)protonation effects, is not captured by the simulations, in which the molecules are represented as strictly zwitterionic.

However, to investigate the difference observed in the simulations between DPPC and DP-DGTS bilayers, in the spirit of our earlier work [28,45], we analyzed the polar headgroup orientation with respect to the membrane normal (Figure 5A). For DPPC, we find that the headgroup is predominantly aligned parallel to the surface, especially at low hydration as previously reported [28]. In contrast, for DP-DGTS the headgroup vector is more perpendicular to the surface, which suggests a more repulsive dipole configuration in line with the more repulsive pressure-distance curve of DP-DGTS. However, a more quantitative analysis of the headgroup dipole moment in the direction normal to the membranes (see inset of Figure 5A and materials and methods) reveals that M_z_ for DGTS is only about half as strong as for PC. Indeed, the electric dipole of DGTS is smaller than that of PC (1 carbon linking the nitrogen to the carboxylate group in DGTS versus 2 carbons and 1 oxygen linking the nitrogen to the phosphate in PC) and the intramolecular charge distribution is different (Figure S7). Thus, the more long-ranged nature of the repulsion observed between DP-DGTS bilayers is not due to the dipole strength or orientation.

**Fig S7.**
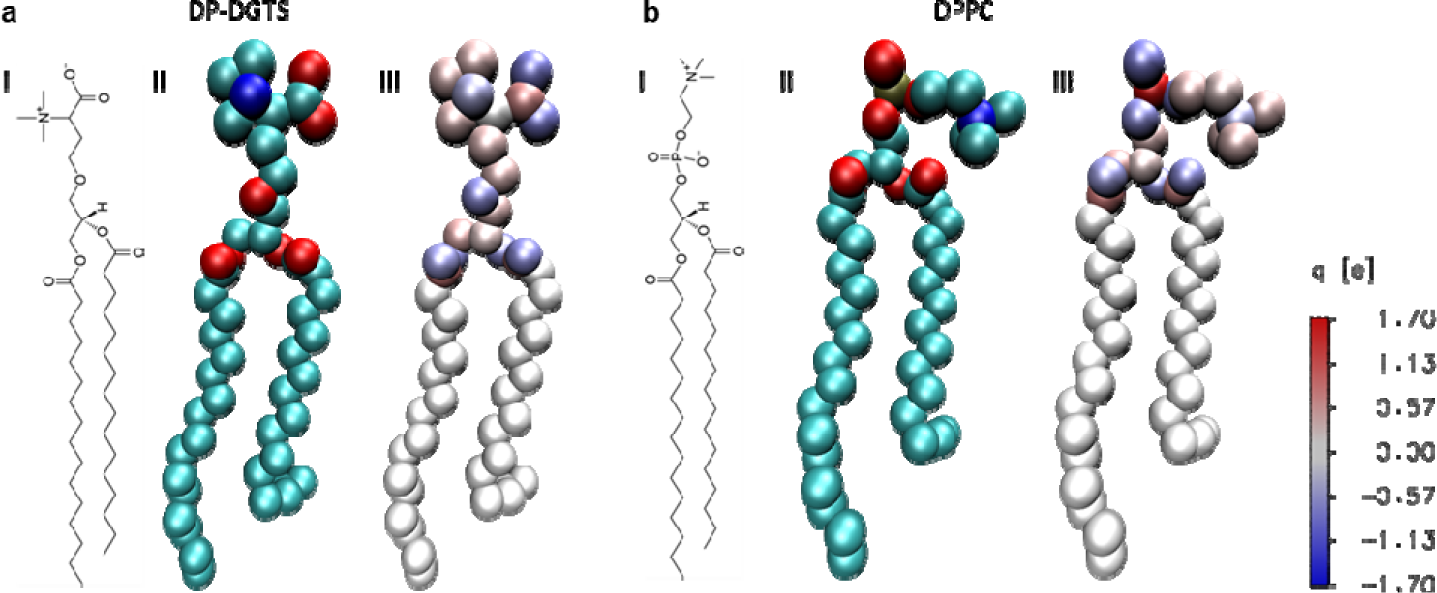
Molecular representation of DP-DGTS (**a**) and DPPC (**b**). (**i**) Shows the chemical structure for both lipids and (**ii**) the united atom representation, where hydrogens are implicitly included on the heavy carbons (green spheres). Red spheres denote oxygen atoms, blue nitrogen atoms and gold phosphor atoms, respectively. (**iii**) Partial charge distribution with color coding on the right. The expected molecular dipole of the zwitterionic headgroups gets largely compensated due to the polarity of the covalent bonds.

**Figure 5:**
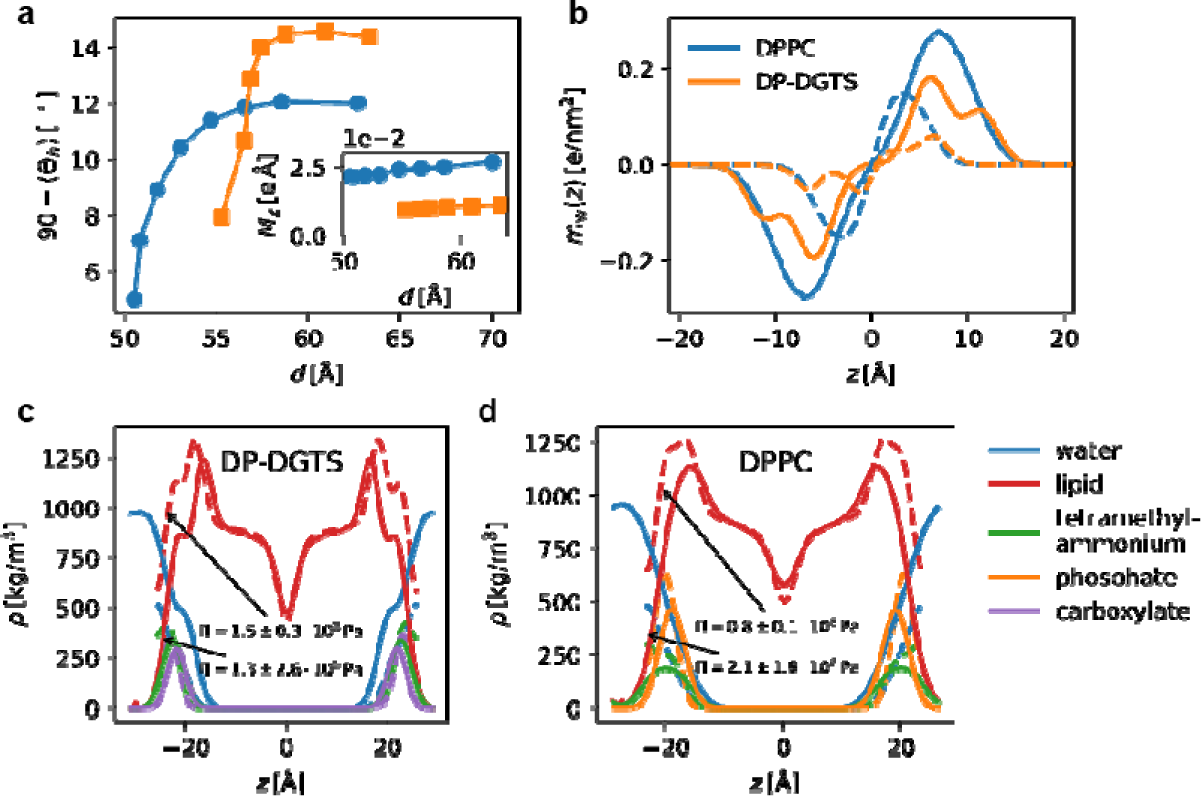
Results of the molecular dynamics simulations. (**a**) Median of the orientation of the DP-DGTS (orange) and DPPC (blue) lipid zwitterionic headgroup vector in fluid phase as function of the repeat distance d. 90°C correspond to an orientation parallel to the membrane plane. The inset represents the lipid dipole moment normal to the surface vs d, indicating that DPPC dipole density is larger than that of DP-DGTS. (**b**) Water dipole density profile versus the distance z along the bilayer normal in fluid phase for DP-DGTS (orange) and DPPC (blue) bilayers, centered here at the middle of the water layer in contrast to panels (**c**) and (**d**), where the bilayer is centered. (**c**) and (**d**) Simulated mass density profiles of the lipids and the water versus the distance z along the bilayer normal for (**c**) DP-DGTS and (**d**) DPPC, centered at the bilayer center of mass. Also shown are the densities of the zwitterionic functional groups of the lipid heads (e.g., phosphate, carboxyl group, trimethylammonium). In **b, c** and **d**, the dash lines represent the profile at low hydration and the solid lines at high hydration.

We then investigated the water orientations profiles (Figure 5B). For DPPC, water is strongly polarized and therefore oriented close to the membrane surfaces and this polarization decays smoothly towards the center of the water layer [28], which is expected from simple continuum modeling [46]. In contrast, for DP-DGTS the water polarization is strongly influenced by the negatively charged carboxylate group inside the polar head region, indicating a strong interaction with water molecules, not present with the phosphate group in DPPC. We therefore extracted the component density profiles of the simulated bilayers to analyze interconnection between water molecules and polar heads (Figure 5C and D). Interestingly, a water layer strongly bound to the polar head of DP-DGTS appears as a shoulder in the profile in Figure 5C, whereas this shoulder is absent in Figure 5D for DPPC bilayers. Therefore, the simulation indicates different interaction with water molecules for DP-DGTS bilayers that could explain partly the observed stronger repulsion between DP-DGTS bilayers. However, the long-ranged distance repulsion not entirely reproduced by molecular dynamics could also be attributed to incomplete charge-neutrality if a small fraction of DP-DGTS molecules deviate from the zwitterionic state.

### Evolution of betaine lipid is influenced by environmental conditions

Our results indicate that the polar heads of DGTS and PC behave differently due to their different chemical structures, which in turn leads to different properties of the formed bilayers. This raises the question of whether the disappearance of betaine lipid through the evolution is linked to these differences. To address this question, phylogenetic analyses were done on the gene involved in DGTS synthesis. DGTS is biosynthesized in a two-step reaction catalyzed either by two enzymes BtaA and BtaB in prokaryotes, or by a bifunctional enzyme Bta1 in eukaryotes enclosing both BtaA and BtaB domains [5,14]. By domain sequence homology, we retrieved Bta1 amino acid sequences from at least 14 families of eukaryotes, bacteria and archaea and performed separate Bayesian phylogenetic analyses on each domain (Figure 6 and Figure S8) because in eukaryotes BtaA and BtaB are arranged in two different configurations [47].

**Figure 6:**
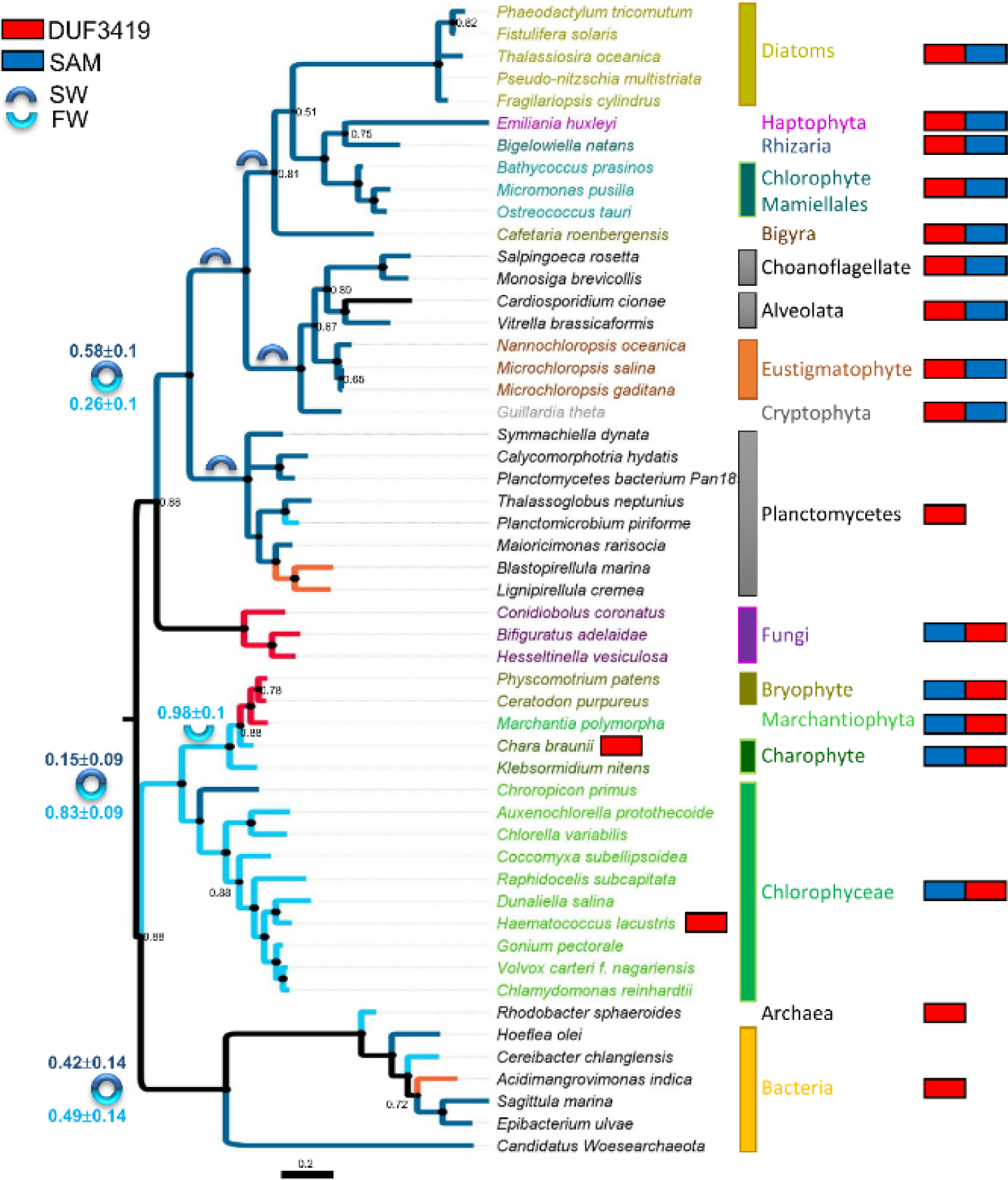
Phylogenetic tree of the BtaA domain of the BTA1 protein from SAR representatives (diatoms, eustigmatophytes), Haptophytes, Cryptophytes, Mamiellales, Chaonoflagellates, alveolates, fungi, bryophytes, charophytes, Chlorophyceae, and bacteria. The tree presented was inferred by Bayesian analysis as described in the Materials and Methods section and the topology overlaps Maximum Likelihood tree topology. Bayesian Posterior Probability (BPP) values below 0.90 are reported at each node. The color of the clades and branches identifies the habitat as validated by bibliographic survey. Dark blue = seawater; light blue = freshwater; red: terrestrial; orange: brackish water. At each internal node, the BPP for the habitat of the ancestor is indicated besides the sign if below 1.00. On the right-hand side the configuration of the protein is reported; red: BtaA domain; blue: BtaB domain.

**Fig. S8:**
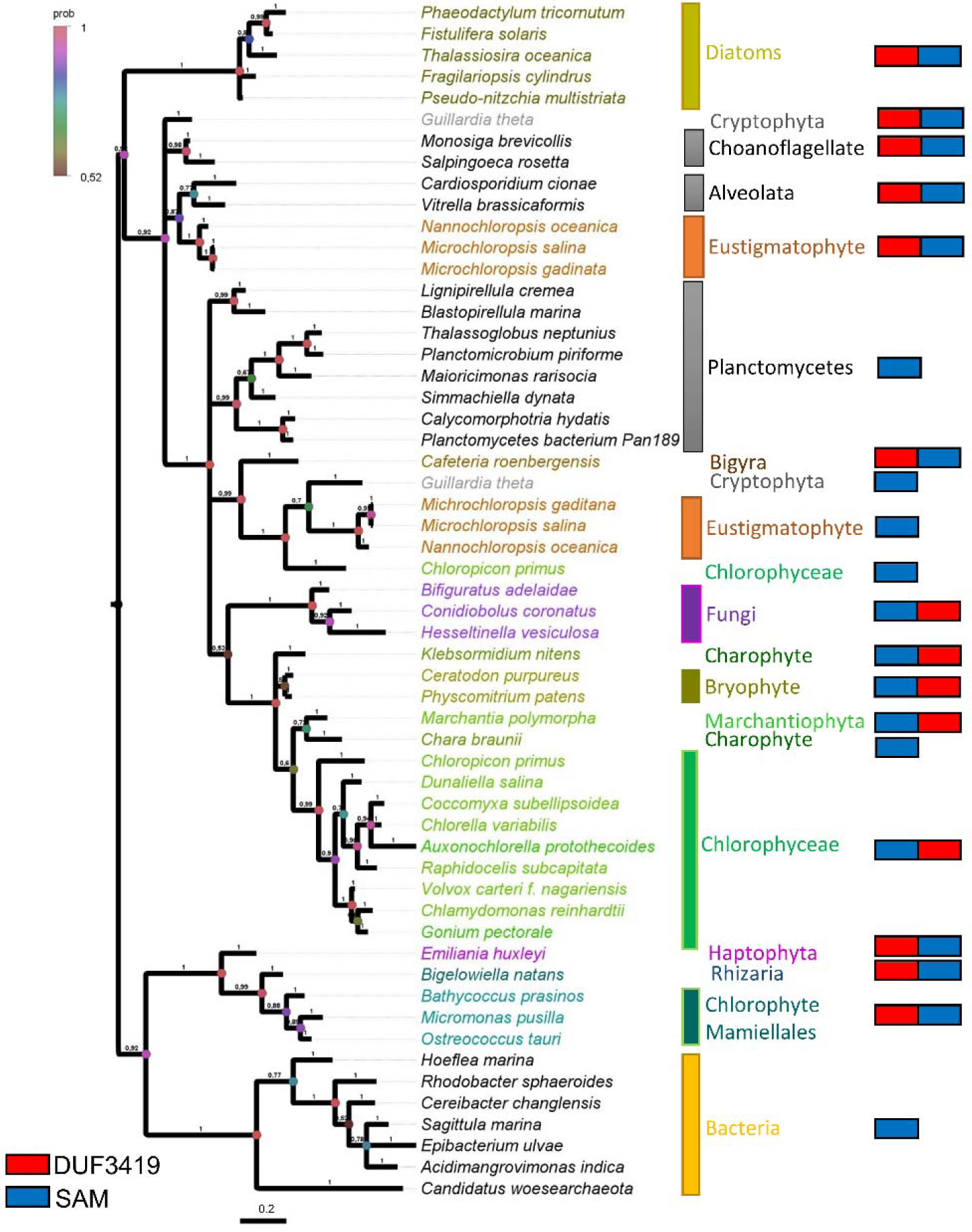
Bayesian phylogenetic tree of the S-adenosyl-L-methionine-dependent methyltransferase domain of the BTA1 protein. Bayesian Posterior Probability values are reported next to the branches. The color of the node circles represents BPP. Color code is reported on the figure. The tree is drawn to scale, with branch lengths measured in the number of substitutions per site. On the right-hand side the configuration of the protein is reported; red: BtaA domain: 3-amino-3-carboxypropyl-transferase domain (Domain of Unknown Function DUF3419); blue: BtaB domain: S-adenosyl-L-methionine-dependent methyltransferase (SAM) domain.

The phylogenetic tree shows a robust separation between marine and freshwater organisms (posterior probability 1.00) and does not corroborate taxonomy [48–50]. Such clustering overlaps with the domain arrangement, i.e. BtaA-BtaB in marine organisms vs BtaB-BtaA in freshwater ones. Interestingly, the planctomycete sequences, a clade of marine bacteria [51], cluster robustly (PP 0.92) at the base of the clade containing all marine protists and algae, whereas Archea and other bacteria form a clade in basal position. Also intriguing is the position of the marine mamiellales (Chlorophyta) such as *Ostreococcus taurii* clustering together with the haptophyte *Emiliania huwleyi* (PP 0.99), and diatoms (PP 0.51) such as *Phaeodactylum tricornutum*, while other terrestrial chlorophytes cluster together in a separate clade.

The habitat was taken into consideration as a character in a bi-partitionned Bayesian analysis and results suggest that either two distinct lateral gene transfers occurred for marine and freshwater organisms, or the two BtaA domains split very early in the evolution (around 1.5 billion years ago) and then evolved separately in marine and freshwater organisms. In this case, the probability that the Last Universal Common Ancestor harbored a BtaA domain similar to the marine or the freshwater organisms is not decisive (0.42 ± 0.14 and 0.49 ± 0.14 respectively). Contrariwise, the Most Recent Common Ancestor of freshwater plants highly likely presented a freshwater BtaA domain (0.83 ± 0.09). Along evolution, habitat switches occur and are detectable as homoplasies (*Chloropicon primus*, a marine species, clusters with liverwort and non-vascularized plants) or synapomorphies (the terrestrial fungi). As a result, the phylogenetic analysis indicates that the evolution of the Bta1 gene might be driven by the physicochemical properties of the environment, such as the osmotic pressure.

## DISCUSSION

Betaine lipids have been reported in the literature for a long time to be a good substitute for phospholipids in extraplastidial membranes, especially under phosphate starvation conditions. We show that DGTS biosynthesis evolution splits between marine and freshwater organisms and does not follow the taxonomy (Figure 6 and Figure S8), suggesting a different selection pressure acting on marine and freshwater environments. Noticeably, marine organisms are naturally rich in very long chain polysunsaturated fatty acids (VLC-PUFAs) - i.e. 20:5 and 22:6 - whereas freshwater organisms contain only medium chain (16 or 18 carbon) PUFAs. A correlation between the marine habitat and the synthesis of VLC-PUFA was suggested [52,53].

We show that DP-DGTS bilayers in fluid phase are on average 6 Å thicker than DPPC bilayers (Figure 3), have higher bending rigidity and a greater tendency than DPPC bilayers to coexist in gel and fluid phase (Figure 1 and 2). Including PUFAs in phospholipid membranes affects the physical properties of the membranes by promoting the fluid phase, increasing disorder and decreasing thickness [54]. *In vivo*, in marine organisms, DGTS and PC do not share the same fatty acid composition. For instance, in *Microchloropsis gaditana*, a eustigmatophyte (stramenopile) marine microalgae, DGTS is richer in 20:5 than PC that contains mainly 16 and 18 carbon fatty acids with 0, 1 or 2 unsaturations [55]. Thus, depleting the PC of VLC-PUFAs could increase the membrane thickness as well as the energy required for deformation, leading to a higher bending rigidity [56]. Therefore, the variation of fatty acid composition observed between PC and DGTS *in vivo* could contribute to have matching physicochemical properties of these two lipids and favor the replacement of one lipid by another depending on the availability of phosphorus in the marine environment [57,58]. This adaptation mechanisms could allow through the evolution marine organisms to produce other forms of betaine lipids such as DGTA and DGCC.

However, freshwater organisms and plants do not produce VLC-PUFA, they mainly contain fatty acids with 16 or 18 carbons. Therefore, there is no possible adaptation of the membrane thickness by changing fatty acid chain length [52]. Freshwater organisms with significant fraction of betaine lipids in their composition usually have little or no PC [59]. Even some algae contain only betaine lipids and no PC such as *Chlamydomonas reinhardtii* whereas some other do not contain betaine lipids but only PC such as *Chlorella* sorokiniana [60,61]. Nonetheless, it was found very recently that in phosphate starvation *Chlorella kessleri* replaced PC by DGTS completely although it was previously thought to be an alga that produces no betaine lipid [62]. Therefore, we could suppose that the inability of freshwater organisms to make PC and DGTS “compatible” through fatty acid chain adjustments forces these organisms to avoid PC/DGTS mixed bilayers. However, these organisms conserve their pathway to synthetize DGTS to maintain the possibility to use betaine lipids to face phosphate starvation by erasing PC.

In land plants, the loss of DGTS synthesis is concomitant to the loss of desiccation tolerance [63]. DGTS was detected in some mosses and ferns but is degraded during dry season [21,64,65] and DGTS synthesis genes are absent in gymnosperms and angiosperms. We showed that DP-DGTS bilayers have more repulsive hydration interactions than DPPC bilayers at short and long-range, possibly due to polar head strong interaction with water molecule and a stronger deviation from the zwitterionic state. In land plants during phosphate starvation, PC is partly replaced by DGDG that has a much less repulsive hydration interaction than PC [28]. Interaction modalities between water and DGTS are possibly deleterious in terrestrial environments. Further investigations are needed to address this question; for instance by producing *Arabidopsis thaliana* containing DGTS and by investigating its properties and membrane architecture during drought.

## MATERIALS AND METHODS

### 1) Lipids

1,2-dipalmitoyl-sn-glycero-3-phosphocholine (16:0/16:0 PC) and 1,2-dipalmitoyl-sn-glycero-3-O-4’-(N,N,N-trimethyl)-homoserine (16:0/16:0 DGTS) were purchased at Avanti Polar Lipids (Alabaster, AL, USA).

### 2) Differential Scanning Calorimetry

Differential scanning calorimetry (DSC) experiments were performed using a Micro DSC III differential scanning calorimeter (Setaram, France). The aluminium capsules containing the samples were filled with 500 µL of 0.1 % and 0.2 % (w/v) solutions of DPPC and DP-DGTS respectively. The lipid suspensions were extruded with a thermostated extrusion kit from Avanti Polar Lipids (Alabaster, AL, USA) by passing the suspensions 11 times through 0.1 µm pore diameter polycarbonate membranes. The two samples were heated and cooled between -4° and 55°C, with a heating rate of 0.5°C/min. Only the last three of four cycles were identical and used in the data analysis, the first one being always different from the following cycles due to the annealing of the vesicles after the extrusion step. The phase transition temperature was defined as the maximum of the endo- and exothermic peaks.

### 3) Neutron experiments

#### 3.1. Neutron diffraction experiments

All neutron diffraction data were collected on the D16 diffractometer of the Institut Laue-Langevin (ILL, Grenoble, France), according to previous works [27]. The wavelength of the neutron beam was λ = 4.47 Å. The experiments were conducted using the BerILL humidity chambers developed at ILL [66] to control in-situ the sample’s hydration and temperature. For both lipids, the temperature in the chamber was maintained at 50 °C during the measurements. The range of humidities investigated was 30 % to 100 % RH by changing the temperature of the water reservoir generating the water vapor. The Ω-scans were collected by steps of 0.05 deg in an Ω-range of -1 to 15 deg. The lamellar periodicity d was calculated using the Bragg’s law as explain in the previous article.

Neutron scattering length density profiles (NSLD) were calculated from the integrated intensities of Bragg peaks corrected for the beam geometry, the neutron absorption (C_abs_), and the Lorentz factor correction (C_Lor_) according to [67], resulting in the corrected discrete structure factor of order n:

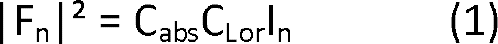

where *I_n_* is the intensity of the Bragg peak at the order *n*.

The corrections are given by:

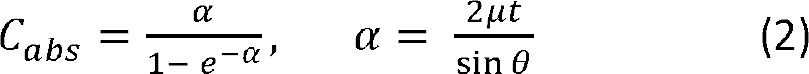

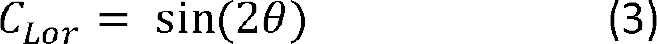

where µ is the absorption coefficient (5 cm^-1^), t is the sample thickness calculated from the deposited amount of dry lipid (0.5 mg) and the sample area (10 cm^2^). It equals 50 µm before hydration, for a deposited amount of 0.5 mg.

The NSLD were constructed with the Fourier transform of the structure factor as follows [68]:

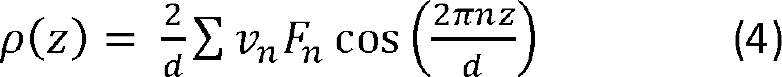

where *z* is the direction perpendicular to the bilayer planes and v_n_ corresponds to the phase of the structure factor of order *n*. According to the literature, at the 8% D_2_O contrast, we tested different hypotheses for the discrete structure factor signs to obtain a centrosymmetric SLD profile with a minimum at z = 0, the bilayer mid-plane, where hydrogen rich methyl groups are located and yield the lowest SLD, and the highest SLD for the polar head regions. The assigned phases that give the best agreement with these constraints were -, -, +, - in agreement with [69] and our molecular dynamics analysis.

We consider here as a definition of the bilayer thickness (*d_b_*) the center-to-center distance between lipid polar heads as obtained from a fit to the two headgroup layer positions in the NSLD profile (as illustrated in Figure S3) [29]. The error associated to *d_b_* is given by the standard deviation between the fit and the calculated NSLD. Finally, the water layer thickness (*d_w_*) is calculated from the known d-spacing d and the bilayer thickness according to:

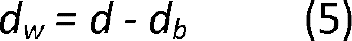

and the error is defined as 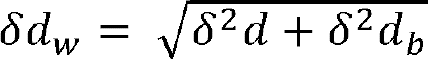.

#### 3.2. Bragg sheet analysis

According to the Discrete Smectic *Hamiltonian* description of interacting multilayers [70] the membrane fluctuation self and cross correlations that give rise to characteristic off-specular scattering are governed by the mechanical properties of the interacting membranes in terms of the membrane bending modulus *κ* and the inter-membrane compression modulus B. As we have shown earlier, the experimentally obtained reciprocal space maps within this framework can be satisfactorily modeled solely based on the underlying mechanical parameters *κ* and B, and on an empirical cut-off parameter termed R [39]. In practice, this procedure relies on the *kinematic approximation* (KA) of wave scattering, because application of the more accurate *distorted-wave Born approximation* [71] would require detailed additional knowledge of the sample structure, which is unavailable. As a consequence, our KA-based treatment, which is only valid wherever the intensity is weak compared to the incident beam, does not correctly capture the specular maximum of the first Bragg sheet, where this condition is typically violated. In the past, we therefore ignored the first Bragg sheet and relied on the second one [36,39]. In line with our more recent work [38], we combine here information from the first two Bragg sheets (see Figure S4): while the Caillé parameter

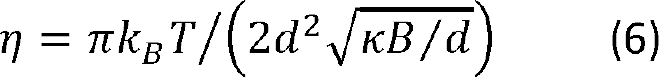

is obtained from the specular/diffuse scattering intensity ratio in the second Bragg sheet, the de Gennes parameter

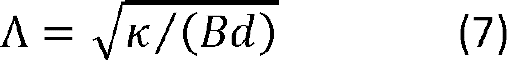

is obtained from the decay of the off-specular intensity in the first Bragg sheet along q_||_, excluding the specular intensity which violates the KA. The best-matching values of *η*, Λ, and R are then determined by their systematic variation in the model until the best agreement with the experimental data is achieved. Finally, the mechanical parameters are obtained by solving Eqs. 6 and 7 for *κ* and B.

#### 3.3. Small angle neutron scattering (SANS)

For the SANS experiments, aqueous solutions of DPPC and DGTS were prepared in pure D_2_O at 10% (w/v) and 3% (w/v), respectively, and then vortexed before transfer into 1 mm path length quartz cells (Hellma, Germany) for the data acquisition. For DPPC, this is well above the swelling limit commonly observed at 40% water for phosphatidylcholines.

SANS data were collected at the D16 cold neutron diffractometer of the Institut Laue-Langevin (ILL, Grenoble, France). D16 uses a monochromator made of nine highly oriented pyrolytic graphite (HOPG) crystals whose orientation can be set to focus the beam vertically in a continuous manner from unfocussed to detector or sample focusing. For the SANS experiments, the D16 instrument was set to pinhole geometry by collimating the beam both horizontally and vertically to produce a symmetrical angular resolution and beam size in the two directions. As described above, the samples were bulk suspensions of multilamellar vesicles prepared in a large excess of pure D_2_O. The sample temperature was changed using a water bath connected to the sample changer. In these conditions the d-spacing measured from the Debye-Sherrer ring observed on the SANS patterns yields the d-spacing at maximum swelling of the lamellar phase with excess water in equilibrium with the membrane stacks.

### 4) Set up of the computer model

The computer model of the hydrated bilayers employs atomistic representations of lipid and water molecules (for simulation snapshots, see Figure S6). Following our previous work [35], we use the assisted freezing method [72] for the construction of fully hydrated membranes in the L_β_ (gel) phase at an initial temperature of *T* = 270 *K*. Initially, 16 lipid molecules in each leaflet are arranged on a hexagonal lattice with random orientation but avoiding strong overlap of the atomic positions. 960 water molecules are inserted randomly in a 3 nm slab, corresponding to *n_W_* = 30 waters per lipid. The system is then carefully relaxed during a 1 ns molecular dynamics simulation forcing the dihedral angles of the lipid tails in gauche transformation as described elsewhere [72]. To enforce a *L*_β_ configuration as shown in Figure S6A, one of the hydrated leaflets including the hydrating waters is then selected and rotated by 180° around the z-axis, after which the system is replicated in the x-y plane resulting in a total of 128 lipid and 3840 water molecules. In all simulation periodic boundary conditions are employed using triclinic unit cells with a lattice angle γ = 60° and the initial area per lipid is fixed to 0.58 nm^2^. This system is then relaxed for 100 ns at 270 K and finally heated up to 300 K using a heating rate of 0.1 K/ns which is sufficiently low to avoid spontaneous melting of the computational model [73]. During the relaxation semi-isotropic pressure coupling is employed using the Berendsen barostat [74] which in the heating step is replaced by anisotropic coupling to allow for full relaxation of the lattice modes.

To construct the corresponding bilayers in the *L*_α_ (fluid) phase, the system is constructed equivalently, but instead of anisotropic pressure coupling a semi-isotropic coupling scheme is employed in the last step at a temperature of 330 K. All scripts employed to set up these systems as well as equilibrated structures and simulation input files are available in https://doi.org/10.18419/darus-2360.

For DPPC, we use the well-established united-atom Berger force-field [75] - which is based on the Optimized Potentials for Liquid Simulations (OPLS) force field with refined parameters for the hydrocarbon tail interactions [76] - and the simple point charge/extended (SPC/E) water model [77]. Although this forcefield is known to be problematic when comparing structural properties from NMR experiments, it reproduces experimental pressure-distance data (Figure S5) [35,45] and the chain melting thermodynamics [73]. Furthermore, comparison with simulations results that use different force fields for lipids and water have shown that the hydration thermodynamics are robust with respect to force field variations [35,78]. Contrary to the original parametrization [75] electrostatic interactions are calculated by the Particle−Mesh−Ewald (PME) method [79,80] using a relative accuracy of 10^-5^ and we checked in Figure S9 systematically the influence of the cutoff for Lennard-Jones interactions by using published experimental values [81–83]. We found that *r_c_*=1.4 nm yields lipid molecular areas and water densities consistent both with the LJ-PME method [84] corresponding to an infinite cutoff and also in agreement with experimental data.

**Fig. S9:**
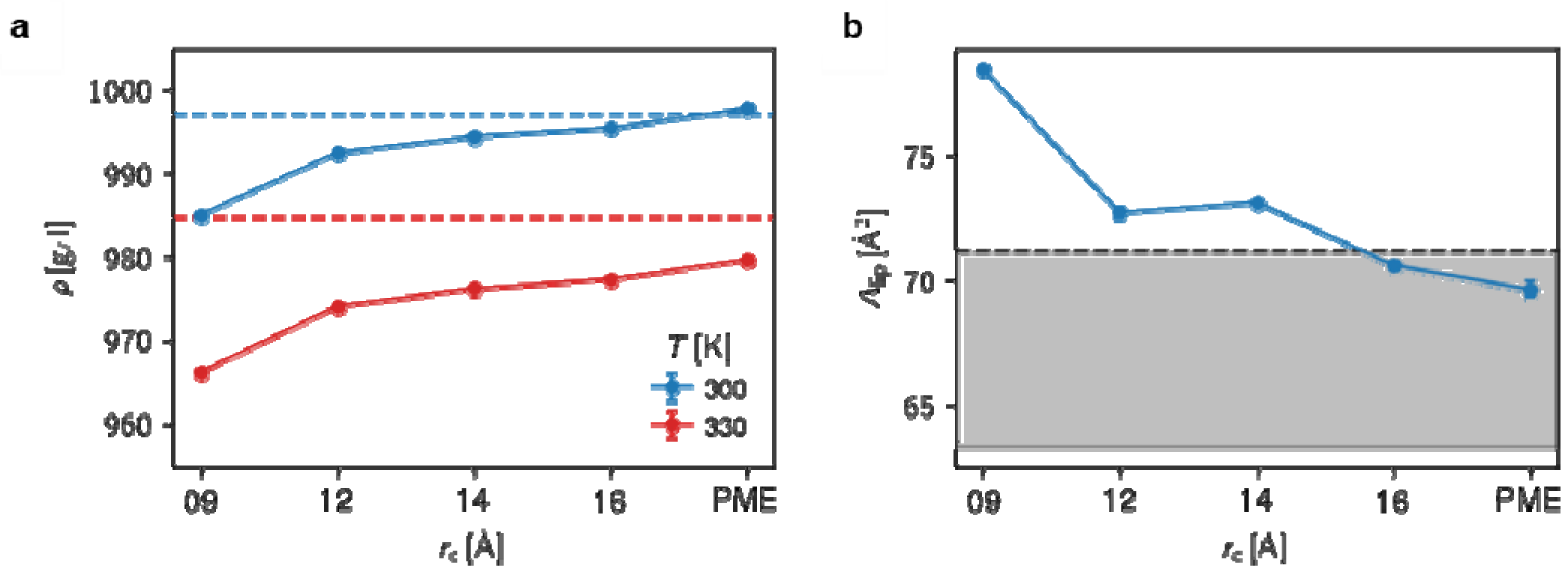
**a.** Influence of the Lennard-Jones cutoff on the bulk water density at 300 K and at 330 K. Vertical dashed lines denote the experimental values [81] **b**. Area per lipid in a DPPC membrane for the different treatment of the Lennard Jones interactions. The gray are denotes the typical values obtained experimentally; the dashed line is obtained by Lis et al. [82], solid line by Petrache et al. [83].

An atomistic representation of DP-DGTS is obtained by adapting the Berger DPPC atomistic topology with refined OPLS parameters for the carboxyl group [85]. In detail, the nitrogen group and the lipid tails are identical and the additional methylene groups are taken from the OPLS force field which results in total charge neutrality on the betaine headgroup. All headgroup bonded parameters stem from OPLS. An initial structure of a DP-DGTS molecule was created using the geometry optimization of the Avogadro simulation package [86]. All the simulation input files are available at https://doi.org/10.18419/darus-2360 and corresponding molecular topologies are displayed in Extended Data Figure 7.

### 5) Computer simulations

All atomistic Molecular Dynamic (MD) simulations are performed using versions 2020 and 2021 of the GROMACS simulation package [87]. All simulations are performed with an integration time step of 1 fs in the canonical constant pressure ensemble as described above. Temperature was maintained at *T* = 300 K for the gel phase and *T* = 330 K for the fluid phase using the canonical velocity rescaling thermostat [88] with a characteristic time of 0.5 ps. The time constant for the pressure coupling is set to 1 ps. Lennard-Jones interactions are truncated and shifted to zero at *r_C_* = 1.4 nm. Water molecules are kept rigid using the SETTLE algorithm [89]. Analysis of the simulations is performed using our freely available MAICoS package (https://www.maicos-analysis.org/) and chemical potentials are evaluated using the Mulistate Bennet Acceptance Ratio method (MBAR) [90] and the toolchain of the Alchemistry project (https://alchemistry.org/)

In the computer simulations, the dehydrating (osmotic) pressure is evaluated using the Gibbs-Duhem relation,

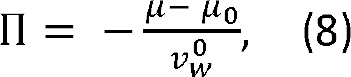

where 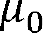 and 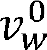 denote the chemical potential and the partial molecular volume, respectively, of pure water in bulk and µ is the chemical potential of water between the bilayers at a given hydration level [45]. Eq. (8) indicates that the chemical potential of water in atomistic MD simulations needs to be evaluated with a precision as high as δµ ∼ 0.01 *k_B_T*. We independently measure the excess and ideal contributions, *µ ^X^* and *µ^id^* = *k_B_T* ln(*ρ_W_*Λ^3^/*m_W_*), where *p_W_* is the water density, Λ its thermal wave length, *N_A_* the Avogadro number and *m_W_* the water molecular mass. *µ ^X^* is evaluated using a Free Energy Perturbation ansatz where a test particle is brought from vacuum to the water using 38 discrete λ states for the MBAR analysis, where first the Lennard Jones interactions are turned on using a soft-core potential approach and then the electrostatic interactions are added. While by definition µ is constant over the simulation volume in thermal equilibrium, its contributions *µ^X^* and *µ^id^* are not due to the inhomogeneous water distribution perpendicular and depend on the coordinate perpendicular to the membrane surface. We thus choose to measure the chemical potential at the center of the water slab between the bilayers and fix the test particle in the plane parallel to the membrane surface using a harmonic potential acting on its center of mass. To overcome potential sampling issues due to the slow dynamics of lipids in the gel phase five independent systems are constructed as described above at *n_W_* = 30 waters per lipid and then step-wise dehydrated by a 100 ns equilibration run. The five equilibrated systems at each *n_W_* are then sampled at each λ state for 100 ns, corresponding to 5 × 38 × 100 ns = 19 µs sampling time per data point shown in Figure 5 and Extended Data Figure 5. For simulations in the fluid phase one system is melted at *T* = 330 K and dehydrated equivalently, where for consistency each ;t state is sampled for 500 ns and analyzed in blocks of 100 ns.

NSLD profiles can be obtained readily from the simulation data by multiplying the united atom density probability with the corresponding scattering length densities, which we took from NIST (https://www.ncnr.nist.gov/resources/activation/). However, this approach corresponds to taking infinitely many terms into account in Eq. (4), which in the experimental analysis is impossible since only few Bragg peaks are present. We thus first calculate the corresponding structure factors |*F_n_*|^2^ via discrete cosine transform and limit to three terms in the reciprocal space, which is the typical number of Bragg peaks present in the experimental analysis. After back transform, this yields NSLD profiles from which the bilayer thickness *d_b_* is evaluated. Note that the phases *v_n_* obtained from the simulations perfectly agree with the experimental analysis. We observe a constant shift in the Π-*d* curves in Extended Data Figure 5 when comparing our simulation results to the experimental data of Δ*d^fluid^* = 3Å for the fluid phase and Δ*d^gel^* = 8Å for the gel phase, which we attribute to the united-atom representation of the lipid tails and consequent underestimation of the chain-chain repulsion between lipid monolayers. This explanation agrees well with the observed larger shift for the gel phase where the tails are more ordered and stretched. Consequently, simulation data for *d* and *d_b_* have been shifted by Δ*d* in Extended Data Figure 5. Density profiles for Figure 5 as well as for the NSLD profiles have been extracted from the simulation trajectory using our open-source analysis framework MAICoS (https://www.maicos-analysis.org/), where in Figure 5 C and D we additionally split into the zwitterionic headgroup parts, the water and the lipid densities. The water polarization profiles in Figure 5B are obtained from the water charge density profiles 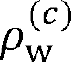 as 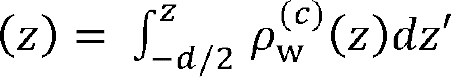, where the lamellar repeat distance *d* equals the size of the simulation box in *z*-direction. Similarly, the dipole moment of the polar heads in the direction perpendicular to the surface is obtained from the polarization densities *m*_l_ of a single membrane leaflet according to 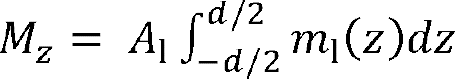, where the subscript l denotes that only the lipid charges are considered in the double integration and *A*_l_ denotes the lateral area per lipid. In detail, for DP-DGTS the tetramethylammonium-carboxylate vector is used, whereas for DPPC the phosphate-tetramethylammonium vector is used. Since by construction of the models, the lipids carry no net charge, this is exactly the dipole moment per unit volume of the membrane.

### 6) Phylogeny

The *Microchloropsis gaditana* locus Naga_100016g36 (annotated as Protein of unknown function DUF3419) was identified as putative betaine lipid synthase by homology with *Nannochloropsis oceanica* BTA1 [47]. The Naga_100016g36 CDS was translated and the amino acid sequence used as a query for a series of BLAST searches in all the publicly available databases. Both tBLASTn and PSI-BLAST [91,92] were performed together with an human-curated process in order to obtain the widest dataset possible and the most robust one. The same was done with all the sequences used by [47]. The final dataset contained amino acid sequences from at least 14 families of eukaryotes, bacteria and archaea. Because in most eukaryotes the two domains of the BTA protein (the 3-amino-3-carboxypropyl-transferase and the S-adenosyl-L-methionine-dependent methyltransferase (SAM) domains) are arranged in two different configurations, two separate alignments were produced. The amino acid sequences of the two domains were compiled in two separate fasta files and aligned using MUSCLE [93] then curated using BMGE (Block Mapping and Gathering with Entropy) software [94] to select the phylogenetic informative regions. For each alignment the substitution model that best fit the data was selected by running the ‘Find Best DNA/Protein Model’ utility implemented in MEGA X (Molecular Evolutionary Genetics Analyses) software [95]. The Le-Gascuel substitution model for Maximum Likelihood (ML) phylogenetic inference method was chosen as the best fitting for both datasets [96]. Non-uniformity of evolutionary rates among sites was modelled using a discrete Gamma distribution (+G). For the SAM domain alignment, a certain fraction of sites were considered to be evolutionarily invariable (+I). The ML phylogenetic analyses were supported by 5000 (bootstrap) pseudoreplicates. Initial trees for the heuristic search were obtained automatically by applying Neighbor Joining (NJ) and BioNJ algorithms to a matrix of pairwise distances estimated using a JTT model, and then selecting the topology with superior log likelihood value. All positions with less than 95% site coverage were eliminated. Trees were drawn to scale with the branch length measured in the number of substitutions per site.

In order to infer both the fusion history and the habitat of the organisms possessing a BTA gene, a Bayesian inference was carried out using MrBayes v3.2.7 [97,98] using a partitioned model [99]. After some trial runs, the conditions for the Bayesian analysis were set up to ensure that the Average standard deviation of split frequencies reached stationarity over the course of the sampling. For each analysis, a total of 1,600,000 generations was implemented, with successive samples separated by 100 generations after an initial “burn in” period of 25 % of the number of samples. The Bayesian posterior probabilities (BPP) were estimated by two independent runs of four Metropolis Coupled chains (MCMCMC). For each dataset, the model selection was done during the analysis by estimating the posterior probabilities of the different models together with their parameters.

## Financial sources

This work was supported by ANR-DFG ANR-18-CE92-0015 and SCHN 1396/2. SB was supported by a joint funding Glyco@Alps / ILL PhD Program, by the French National Research Agency in the framework of the “Investissements d’avenir” program Glyco@Alps (ANR-15-IDEX-02), “Origin Of Life” (ANR-17-EURE-0003), “Oceanomics” (ANR-11-BTBR-0008) and the Labex GRAL, financed within the University Grenoble Alpes graduate school (Ecoles Universitaires de Recherche) CBH-EUR-GS (ANR-17-EURE-0003). AS acknowledges funding from the DFG under Germany’s Excellence Strategy – EXC 2075 – 390740016. Computing time was provided by the state of Baden-Württemberg through bwHPC.

## Data Availability Statement

The data that support the findings of this study are openly available at https://doi.ill.fr/10.5291/ILL-DATA.8-02-220, https://doi.ill.fr/10.5291/ILL-DATA.TEST-3119 and at https://doi.org/10.18419/darus-2360.

## Author Contributions

JJ, BD and ES conceived and designed the research; SB and BD performed the DSC and neutron diffraction experiments, TM performed the SAXS experiment, AS performed the molecular dynamic simulations, OB and AA did the phylogeny analysis. SB, BD, JJ, AS and ES analyzed and interpreted the data. All authors were involved in drafting and revising the manuscript.

