## Supplementary figures and images for "The possible role of lipid bilayer properties in the evolutionary disappearance of betaine lipids in seed plants"

### Figure S1

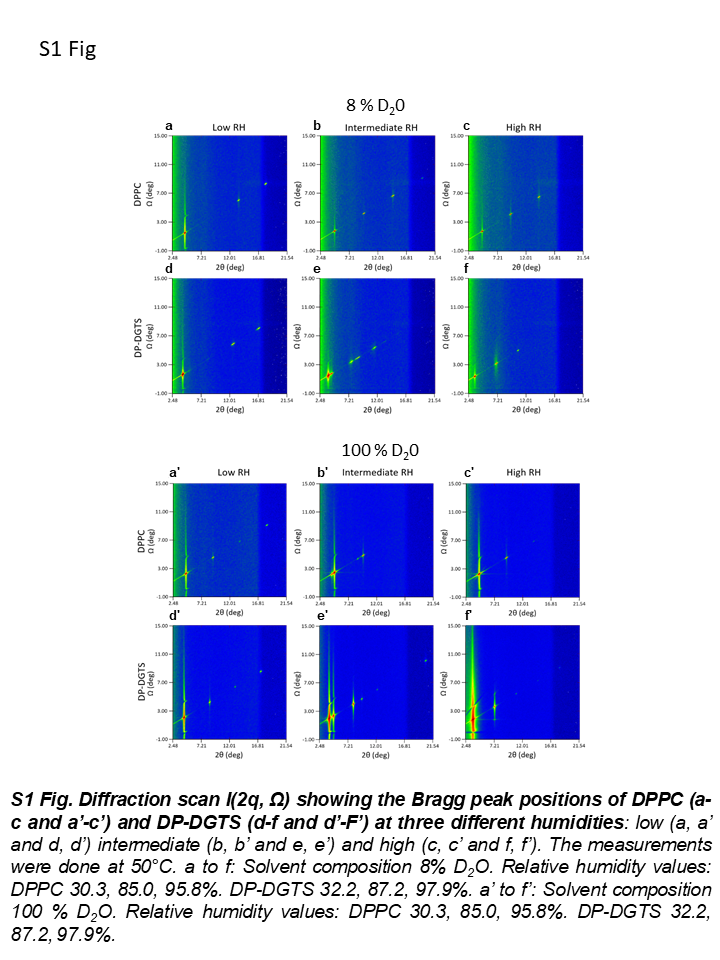

### Figure S2

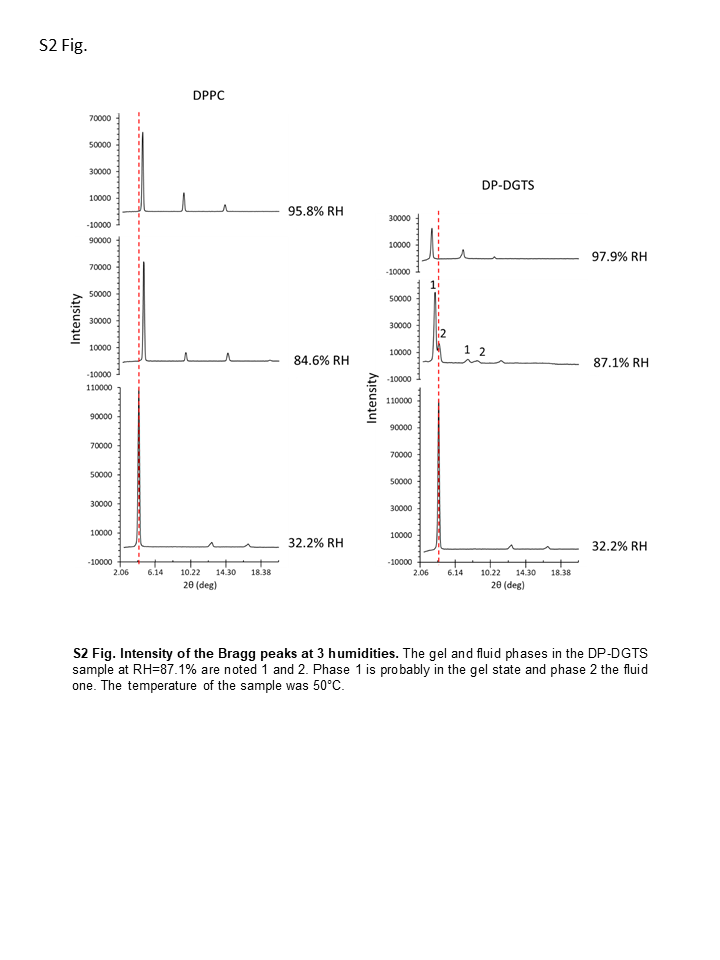

### Figure S3

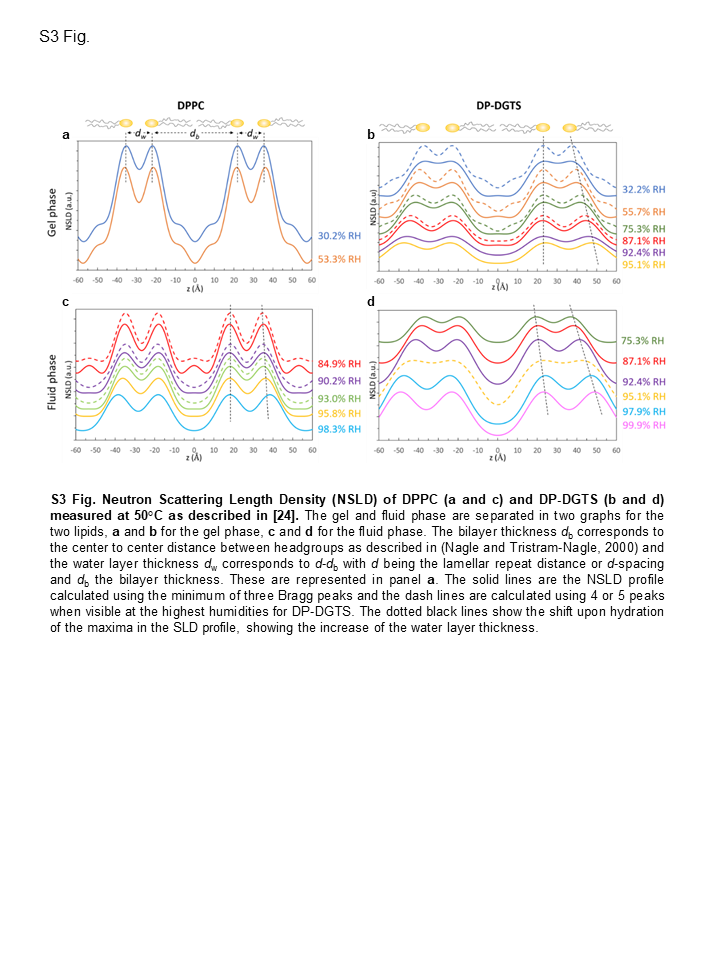

### Figure S4

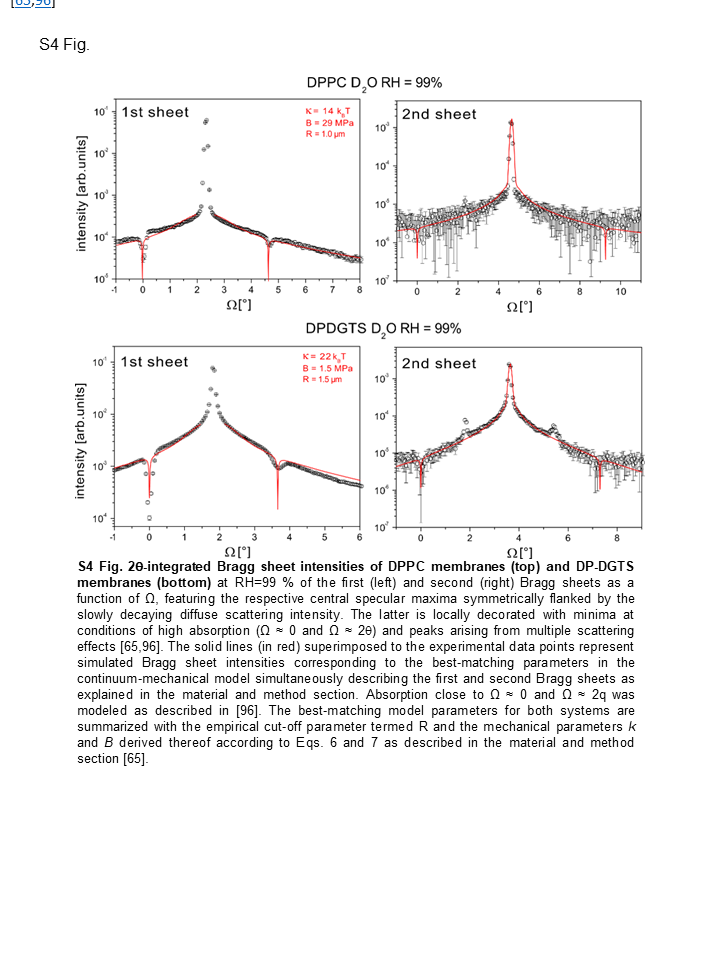

### Figure S5

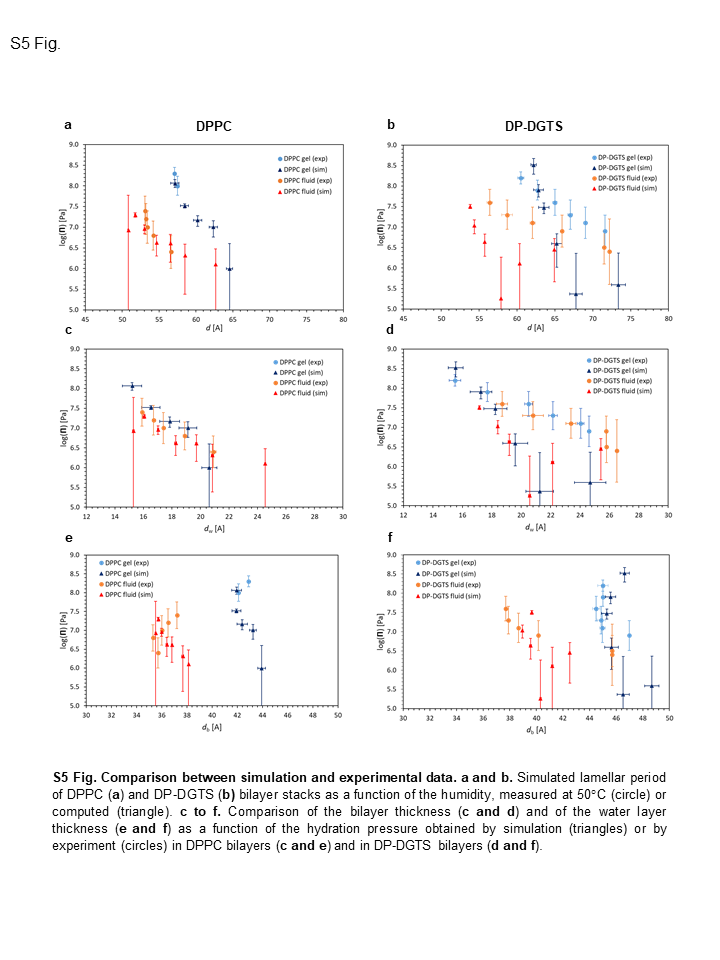

### Figure S6

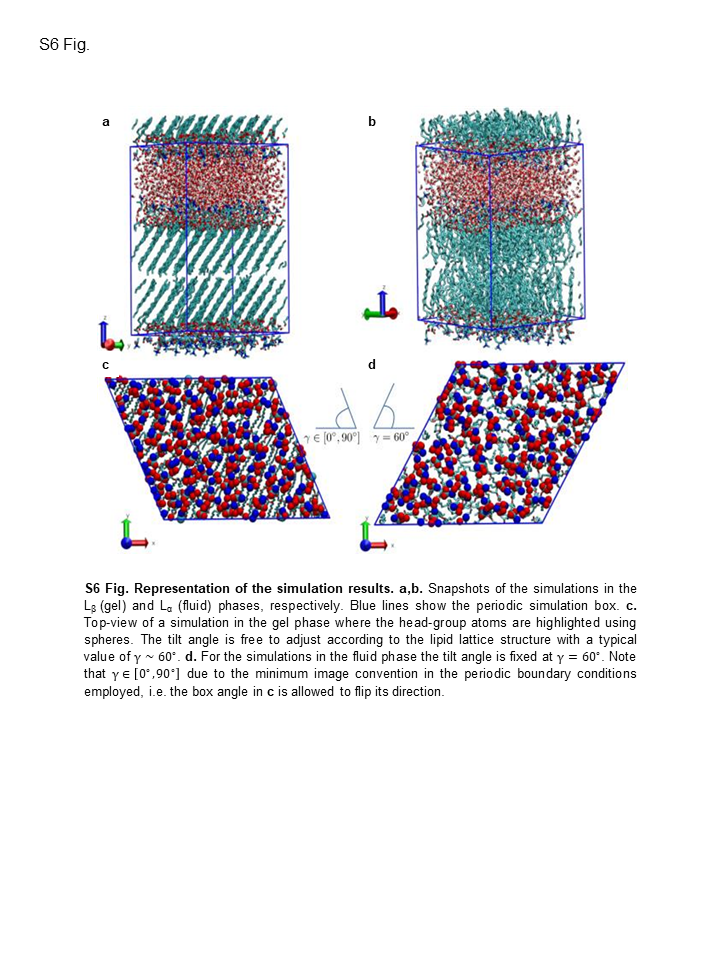

### Figure S7

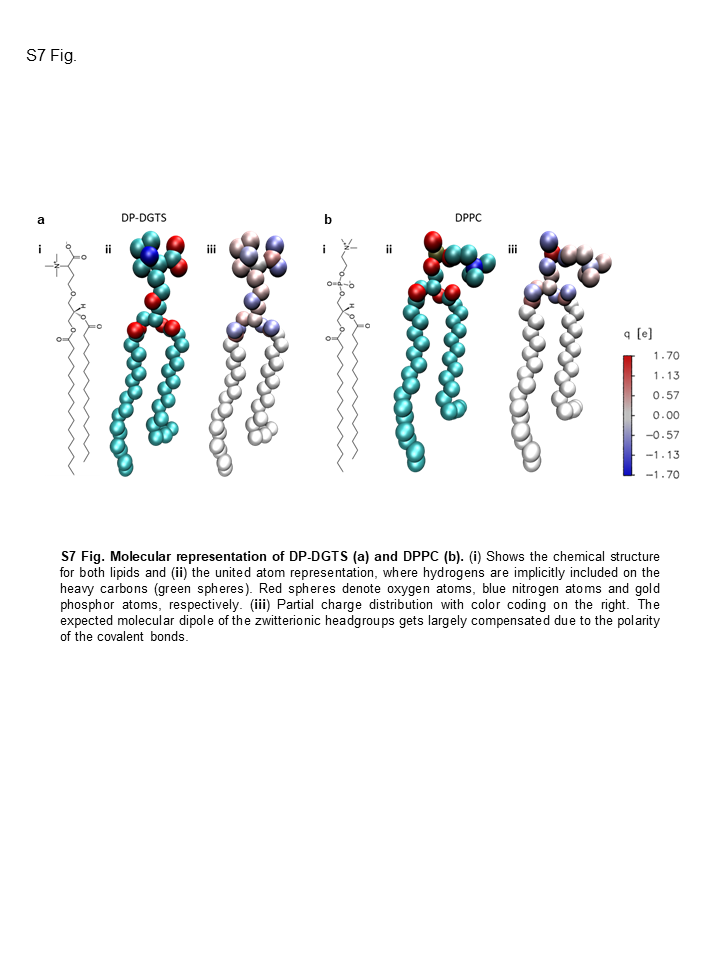

### Figure S8

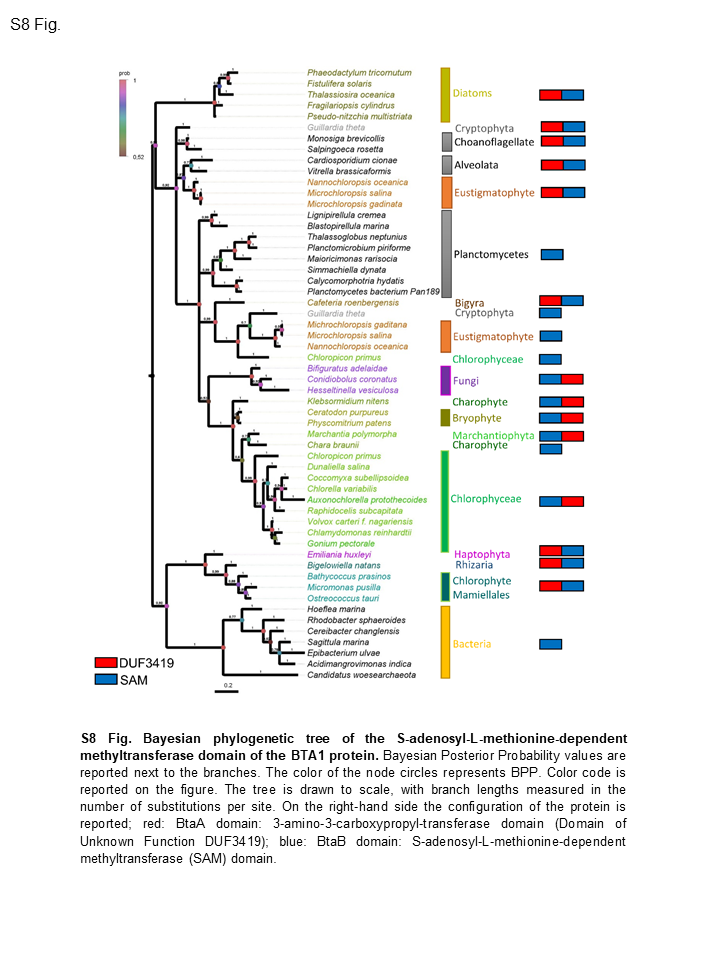

### Figure S9

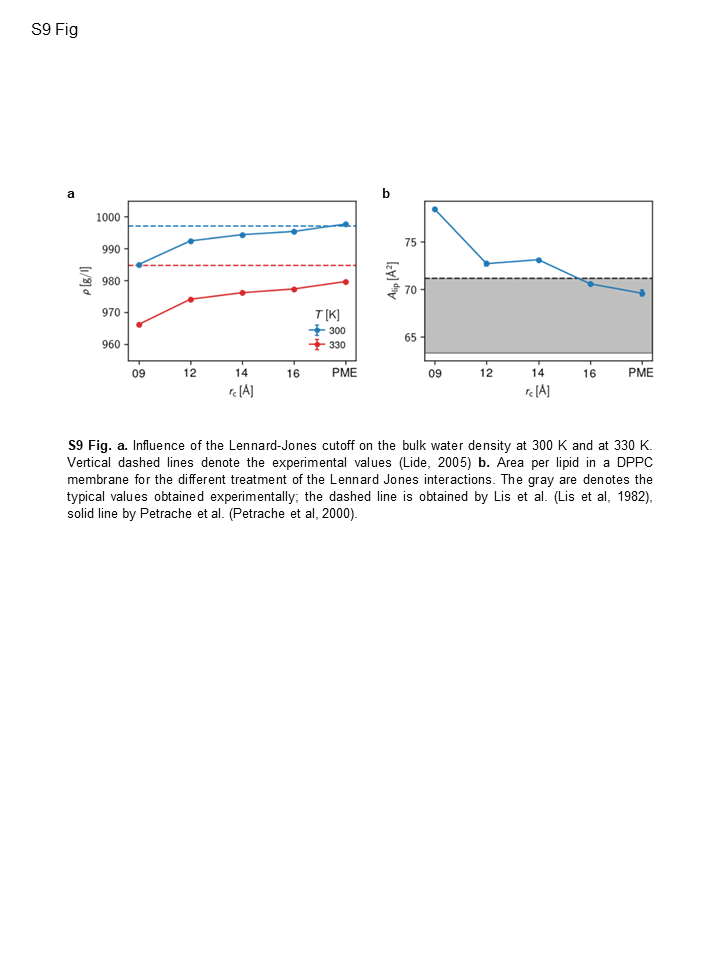
